# Third-order motifs are sufficient to fully and uniquely characterize spatiotemporal neural network activity

**DOI:** 10.1101/2021.08.16.456546

**Authors:** Sarita S. Deshpande, Graham A. Smith, Wim van Drongelen

**Affiliations:** Medical Scientist Training Program; Committee on Neurobiology; Section of Pediatric Neurology; Committee on Computational Neuroscience, The University of Chicago, Chicago, IL 60637, USA

**Keywords:** network analysis, triple correlation, motif class spectrum, third-order correlation, bispectrum, triple correlation uniqueness, triplet motif

## Abstract

Neuroscientific analyses balance between capturing the brain’s complexity and expressing that complexity in meaningful and understandable ways. Here we present a novel approach that fully characterizes neural network activity and does so by uniquely transforming raw signals into easily interpretable and biologically relevant metrics of network behavior. We first prove that third-order, or triple, correlation describes network activity in its entirety using the triple correlation uniqueness (TCU) theorem. Triple correlation quantifies the relationships among three events separated by spatial and temporal lags, which are triplet motifs. Classifying these motifs by their event sequencing leads to fourteen qualitatively distinct motif classes that embody well-studied network behaviors such as synchrony, feedback, feedforward, convergence, and divergence. Within these motif classes, the summed triple correlations provide novel metrics of network behavior, as well as being inclusive of commonly used analyses. We demonstrate the power of this approach on a range of networks with increasingly obscured signals, from ideal noiseless simulations to noisy experimental data. This approach can be easily applied to any recording modality, so existing neural datasets are ripe for reanalysis. Triple correlation is an accessible signal processing tool with a solid theoretical foundation capable of revealing previously elusive information within recordings of neural networks.

## 1 Introduction

Meaningfully capturing the complexity of neural networks is a daunting task: a range of different approaches are currently in use to quantify network behavior [1, 2, 3, 4, 5, 6, 7, 8]. Intuitively, we assume that completely characterizing this complexity should require correlations of such high-order as to be incomprehensible, perhaps even unenumerable [9]. However here we show that *third-order* correlations fully characterize even the most complex neural recordings.

This full characterization arises from the unique correspondence between a dataset and its triple correlation, per the triple correlation uniqueness (TCU) theorem. Introduced decades ago in optical sciences [10], this theorem states that any finite image has unique triple correlation. The TCU theorem has since languished^1^, but we hope to bring triple correlation to the forefront of data analysis with one simple observation: any finite dataset can be interpreted as an “image.” In neuroscience, this encompasses any completed recording of neural activity: functional magnetic resonance imaging, local field potential, single-unit electrophysiology, multi-electrode array electrophysiology, voltage-sensitive dye imaging, etc. Beyond these, the TCU theorem applies to any finite dataset in any field.

Here, we bring the TCU theorem into the heart of neuroscience with a focus on analysis of spike rasters as the representation of neuronal population activity. We propose that the activity patterns of a neural network can be ideally characterized by its triple correlation, *c*_3_. We begin by defining *c*_3_, which comprises triplet motifs, the relationships among three events. We briefly summarize the proof of the uniqueness of *c*_3_ and explore the implications of this uniqueness for neural data. We then classify the triplet motifs by event sequencing and neuron recruitment to derive a natural summary of *c*_3_ in fourteen motif classes. This summary falls out naturally and maps onto network properties of both computational and biological interest, including synchrony, feedback, feedforward, convergence, and divergence. We illustrate the utility of that simple summary with some straightforward simulations and apply our analysis to experimental recordings. In sum, we present a novel, theory-based approach to quantifying network activity.

## 2 Methods

We worked with a multi-neuron raster of spikes (Fig. 1A) to outline our approach, though the analysis works the same for any finite dataset (see Supporting Materials).

**Fig. 1.**
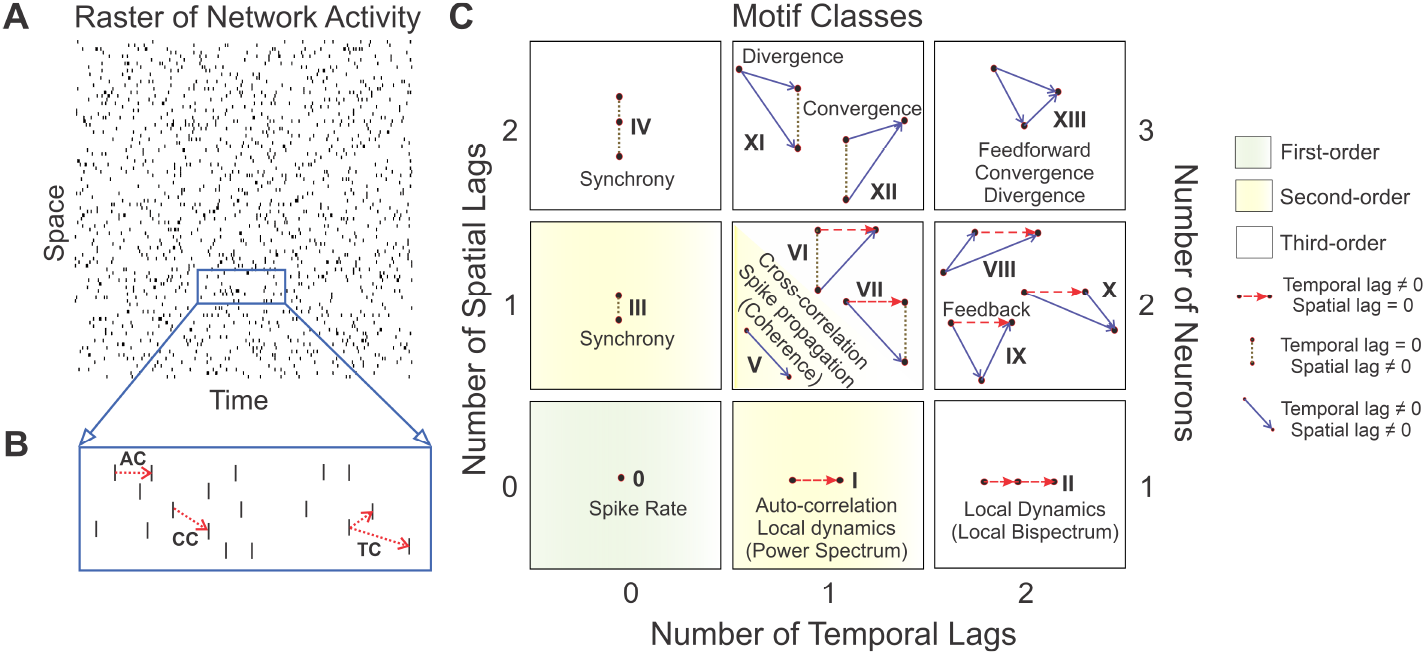
Application of the fourteen motif classes to spike rasters. For a given raster (Panel A), second-order correlations can relate activity within a neuron (auto-correlation, AC; Panel B) or between neurons (cross-correlation, CC; Panel B). Triple correlation (TC; Panel B) relates three bins, separated by up to two temporal and two spatial lags. Three particular spike bins constitute a triplet motif (e.g. as shown in Panel B). We classify these motifs into fourteen motif classes by the motif’s spike sequence (Panel C; see Online Methods for a complete derivation). Dot: a single spike bin. Horizontal red dashed arrow: intraneuronal spike bins, i.e. space lag = 0, e.g. I and II. Vertical stippled line: synchronous spike bins, i.e. time lag = 0, e.g. III and IV. Solid blue arrow: interneuronal spike bins (e.g. V). These 14 motif classes can also embody well-known neuronal processing properties (such as synchrony, feedback, convergence, divergence, and feedforward) in both the time (listed adjacent to the motif class) and frequency domains (listed within parentheses). First-(0) and second-order (I, III, and V) motif classes are highlighted in green and yellow, respectively. The second-order motif classes I, III, and V are constituent motif classes that comprise the third-order motifs. The remaining ones are third-order motif classes.

### 2.1 Defining triple correlation

Unlike pairwise correlation (a function of one lag between two spikes), triple correlation characterizes three-way interactions as a function of **two** lags among three spikes (Fig. 1B). We will call C_3_ the triple correlation transform, which transforms a spatiotemporal raster into its triple correlation. So for a spatiotemporal raster, *r*(*n, t*) (*n*, space; *t*, time), the triple correlation *c*_3_ is a function of four variables: C_3_[*r*] = *c*_3_(*n*_1_, *t*_1_, *n*_2_, *t*_2_) = ⟨*r*(*n, t*)*r*(*n* + *n*_1_, *t* + *t*_1_)*r*(*n* + *n*_2_, *t* + *t*_2_)⟩_*n,t*_. The operation ⟨…⟩_*n,t*_ computes the average over all bins (*n, t*) in the raster. The argument … is 1 when (*n, t*) is part of a spiking triplet with lags (*n*_1_, *t*_1_, *n*_2_, *t*_2_), and 0 otherwise. In practice, we only calculate the triple correlation up to a certain maximum spatiotemporal lag which we determine based on experimental and computational considerations on a per-experiment basis (noted in each figure). In most cases calculating the full *c*_3_ would be a needless computational expense as usually the triple correlations with longer lags become more likely to be dominated by chance. See “Computing the triple correlation” in Online Methods for implementation details.

### 2.2 Overview of uniqueness proof

To prove uniqueness, we apply the Triple Correlation Uniqueness (TCU) theorem, originally applied in optical sciences [10, 13, 14]. It states that if we have two images *x* and *y* with equal triple correlations (C_3_[*x*] = C_3_[*y*]), then the images themselves must be equal (*x* = *y*). The TCU theorem applies to any finite bounded dataset, including continuous signals and multi-dimensional data—a fact that can be understood by interpreting any dataset as an image— so we apply it to a spike raster. We begin with two rasters *x* and *y* whose triple correlations are equal. This implies that the triple correlations’ Fourier transforms (bispectra) are equal, which we can write as equality between the product of three characteristic functions. Characteristic functions are the Fourier transform of probability distributions and are well-known in statistics to have certain nice properties [16]. Using those properties, we perform simple algebraic manipulations of that equality to derive that the Fourier transform of *x* equals the Fourier transform of *y* times an exponential function, F[*x*](*σ*) = F[*y*](*σ*)*e*^*jaσ*^, where *σ* is frequency, *j* is the imaginary number, and *a* is some constant. This means that raster *x* equals raster *y* translated by some constant *a*. Thus, rasters with equal triple correlations are themselves equal up to translation See the Online Methods for the full proof, both in the two dimensions of a typical raster and in the fully general *N* dimensions. As a consequence of this result, spiking activity can in principle be recovered in its entirety from triple correlation (see “Reconstruction Algorithms for Finite Images” in [13]).

### 2.3 Summarizing triple correlation with motif classes

In order to better interpret triple correlation, which is inherently highdimensional, we summarized it along lines meaningful to the underlying neural dynamics, combining the motifs into qualitatively distinct motif classes, *M*_*i*_. To do this, we asked what differences between three-spike motifs constitute fundamental differences. First, we grouped motifs according to whether their lags are zero or non-zero, producing 2^4^ = 16 groups for the four spatio-temporal lags (in Table S2’s numbering, the first number). Next we distinguished within these groups according to the signs of the lags, which expanded those 16 groupings into 169 lag-sign motifs (proven in the Online Methods section “The number of lag-sign motifs;” enumerated in Table S2). These 169 lag-sign motifs enumerate all possible shapes of the motifs where the identity of each node matters, and where space is ordered. However, for our purposes many of these shapes are functionally the same. For example, we only distinguished between zero and non-zero spatial lags: zero meaning “intra-neuronal” and non-zero meaning “inter-neuronal.” Further, node identity is irrelevant. Using these considerations, we grouped together these 169 lag-sign motifs according to those invariant under reflecting over the horizontal axis and node identity permutation (compare “Configurations” in Table S2). As a result, we found fourteen motif classes (Fig. 1C; proven in the Online Methods section “The number of motif classes.”).

### 2.4 Controlling for expected contributions

When using motif classes to quantify network patterns, it is essential to distinguish between the occurrence of these motif classes due to underlying network behavior versus due to chance. Based on simple combinatorics, we expect contributions for each motif class to differ by orders of magnitude. For example, motif classes I and III each have only one varying lag, whereas motif class XIII has four. So, if *λ* is the maximum lag between spikes in a motif, the expected contributions of motif classes I and III are *O*(*λ*), while the expected contribution of motif class XIII is *O*(*λ*^4^). Intuitively, we also expect the motif class contributions to vary depending on the number of spikes required in the motif: while there are more motifs in motif class XIII, the chances of any one of them occurring is lower than motif class I because a motif needs three spikes for motif class XIII but only two for motif class I. So given a fixed spike rate *p*, we calculated the theoretically expected motif class contributions under the simplifying assumption that every neuron is a Poisson process with rate *p* per bin: 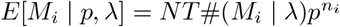 where *NT* is the size of the raster, #(*M*_*i*_ | *λ*) is the number of motifs given maximum lags *λ*, and *n*_*i*_ is the number of events in a motif in motif class *i*. (See Online Methods “Expected contributions per motif class” for case-by-case calculation)

We note that the higher-motifs are composed of lower-order motifs, e.g. looking at motif class X in Fig. 1C, it is constructed of two kinds of arrows: those of motif classes I and V. We say that motif classes I and V are constituent motif classes of motif class X (see also Fig. 4 for more explicitly deconstructed examples). We want to control also for this dependency. By computing the expected relationship between the contribution of a higher-order motif class and the contributions of its lower-order constituent motif classes, we derived *E*_*c*_[*M*_*i*_] which is the expected motif-class contributions given a spike rate and controlling for lower-order constituent motif-class contributions (constituent-controlled expectation). For lower order motifs, this is the same (e.g. *E*_*c*_[*M*_*I*_] = *E*[*M*_*I*_]), but for higher order motifs this reflects the expected contribution in a raster of Poisson processes with the observed contributions of its constituent motifs, i.e. *E*_*c*_[*M*_*X*_] ≈ *E*[*M*_*X*_ | *M*_*I*_, *M*_*V*_]. We report *M*_*i*_*/E*_*c*_[*M*_*i*_] − 1 so that positive values indicate higher contribution than expected, negative indicate less, and zero indicates contributions in line with those expected due to noise and the observed lower-order constituent motif-class contributions.

## 3 Results

Our principal goal is to bring triple correlation into general usage, with the triple correlation uniqueness theorem providing theoretical foundation for its use. Since triple correlation is entirely untested in the field, we approached it as we would any new tool: we showed that it works in the simplest possible case, before adding noise and testing it across the whole gamut of possible patterns. After these checks, we applied triple correlation to open-source real-world data.

### 3.1 Application of triple correlation summary

For a simple example to test this approach, we simulated a network spike raster with synchronous, periodic firing at a frequency of *f* = 0.08 arbitrary units (Fig. 2A). From this raster, we determined contributions of all motif classes by summing the raster’s triple correlation across all motifs in each motif class (*M* = {*M*_*i*_}_*i*∈0:XIII_, where *M*_*i*_ = ∑ *c*_3_(*m*) for motifs *m* in motif class *i*). We also simulated rate-matched pure-noise rasters (Fig. 2B) as surrogates and calculated their motif-class contributions (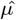 where *μ* = {*E*[*M*_*i*_]}_*i*∈0:XIII_). These surrogates were simulated Poisson processes with identical firing rates for every neuron. In addition we calculated a theoretical value, the constituent-controlled expectation (*μ*_*c*_ = {*E*_*c*_[*M*_*i*_]}_*i*∈0:XIII_; see Methods). We plotted all three quantities for each motif class (Fig. 2C). These quantities on their own are difficult to read: any difference between motif classes is completely overshadowed by the inevitable combinatorial differences in the number of triplet motifs per motif class. To account for these expected combinatorics and effectively report the contribution of each motif class, we calculated the constituent-controlled ratio (*M/μ*_*c*_ − 1; Fig. 2D), which highlights the deviation of each motif-class contribution from that expected due to noise and lower-order constituent motif class contributions. To demonstrate that the contributions *m*_*i*_ from surrogate rasters do not differ much from the constituent-controlled expectation *E*_*c*_[*M*_*i*_], we plotted the estimated-to-constituent-controlled ratio 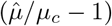 for all *n* = 100 surrogate rasters (Fig. 2E).

**Fig. 2.**
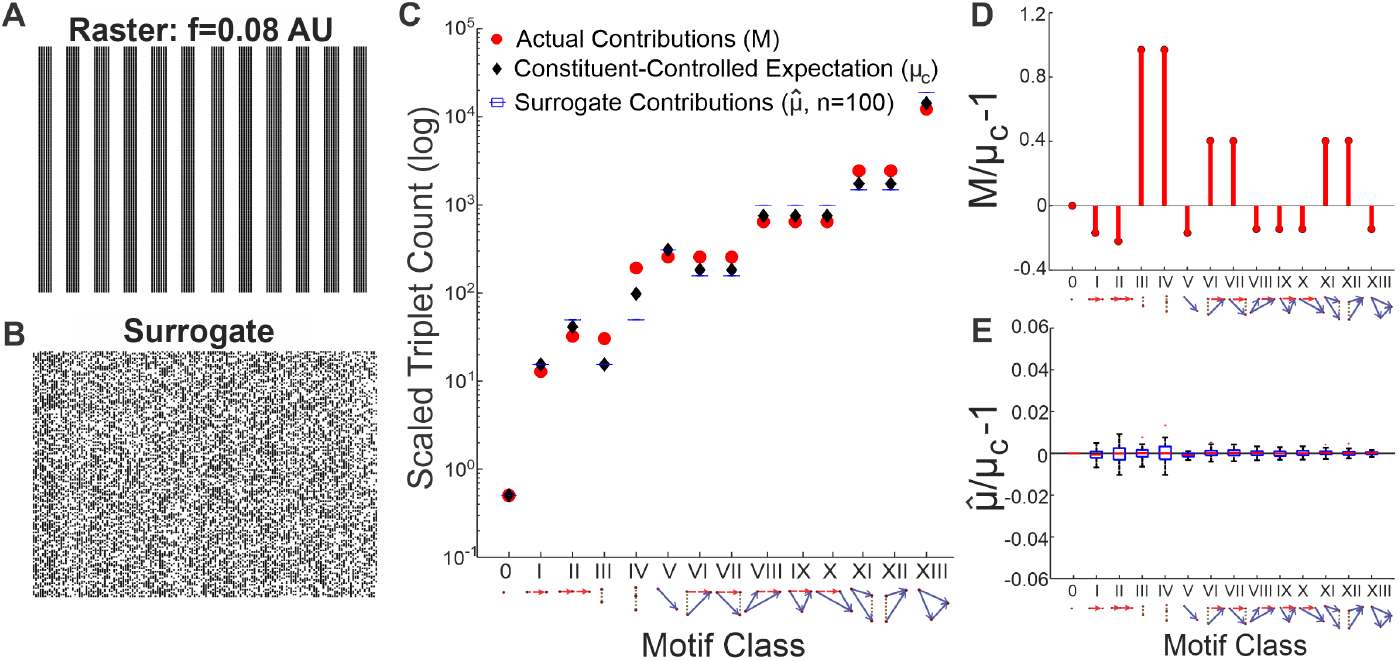
Motif-class contributions to the raster’s triple correlation. A) A 150 *×* 150 spike raster generated by thresholding an 0.08 AU frequency sine wave. B) Surrogate raster generated by randomly shuffling the periodic raster in panel A. C) Various motif-class summary metrics calculated from a triple correlation using lags up to 20 bins in time and space: the actual contributions per motif class (M, red circles); the constituent-controlled theoretically expected contributions (*μ*_*c*_, black diamond) conditioning on spike rate and controlling for the observed contributions of lower-order motifs; surrogate contributions (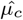, blue boxplots visible as horizontal lines due to relatively small variance) the average motif-class contributions across *n* = 100 shuffled surrogate rasters. D) The constituent-controlled ratios (*M/μ*_*c*_ *−* 1) per motif class. *M/μ*_*c*_ *−* 1 for purely synchronous motif classes III and IV are highest; motif classes VI, VII, XI, and XII (which all consist an element of synchrony) also show positive *M/μ*_*c*_ *−* 1 values. Note that motif class 0 always has zero signal because motif class 0 is the spike rate, which is controlled for by both N and T. E) The estimated-to-constituent-controlled ratios 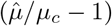 for 100 noise simulations fluctuate around 0 for all motif classes.

We see that our summary of the triple correlation reflects the simple underlying structure: motif-classes III and IV, which correspond purely to neural synchrony, are highest above the expected contributions. Motif-classes VI, VII, XI, and XII, which each includes a synchronous component, are also above chance expectations (Fig. 2D). Thus as a single facet of our analysis, we are able to detect not only second-order synchrony (motif class III), which is considered important in neuroscience research [17, 18, 19], but also third-order synchrony (motif class IV). We recognize that this is an ideal case, in which the synchronicity is also visibly apparent from the raster itself. We will now proceed to show that this particular detection works in the face of noise, and that all the other facets of our analysis (i.e. the other motif classes) also work.

### 3.2 Noise is no obstacle

We explored the effects of increasing noise on detecting network structure using faster synchronous firing with added uniform noise input (*f* = 0.12 AU, SNR = 0 dB; Fig. 3A). Similar to the results of those with slower synchronous firing (Fig. 2D), motif classes III, IV, VI, VII, XI, and XII (those with elements of synchrony) are found more often than chance (Fig. 3B). We increased noise (SNR = −9*dB*; Fig. 3C), resulting in lower magnitude signals across all motif classes, yet still detecting synchronous signals (Fig. 3D). In this case, the underlying network structure is still overtly present in the raster. We then increased the noise such that the synchronous network structure in the raster is not overtly present (SNR = −17*dB*;Fig. 3E), and found that while all signals move even closer to 0, those with synchrony are still elevated (Fig. 3F). Thus, triple correlation reflects underlying structure in the face of added noise. Finally, we show that in an extremely noisy raster (SNR = −40*dB*; Fig. 3G), the synchronous signals are no longer detected as all motif classes now show even lower magnitude signals, approaching 0 (Fig. 3H). In this overwhelmingly noisy simulation, the motif-class spectrum is dominated by chance.

**Fig. 3.**
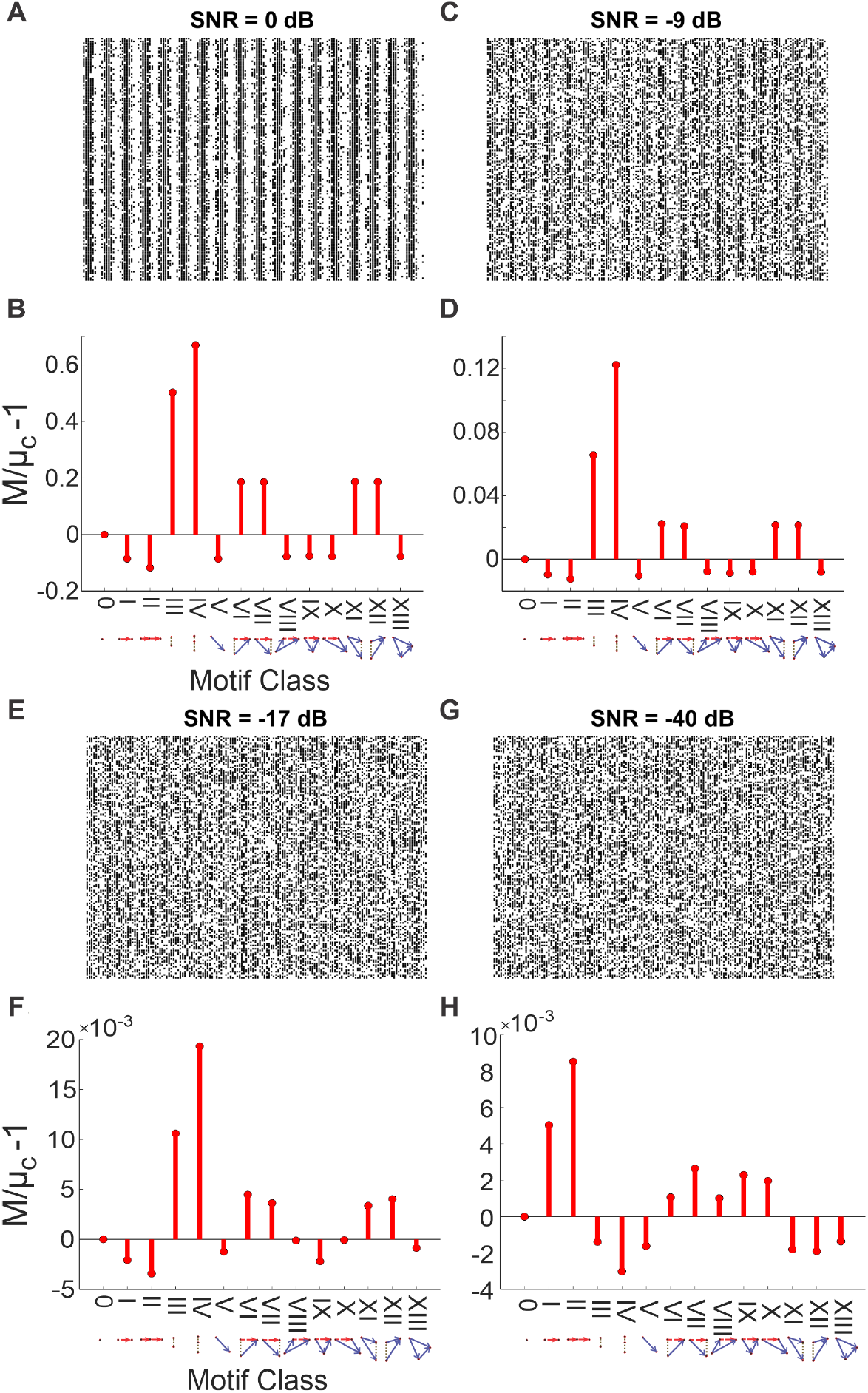
Detecting synchrony amidst increasing noise. A) The 150 *×* 150 spike raster plot generated by thresholding a 0.12 AU frequency sine wave with added noise. The noise consists of uniform noise scaled to give the desired signal-to-noise ratio (SNR = 0 dB). B) The motif-class contributions (*M*) of the above raster relative to chance (*M/μ*_*c*_ *−* 1). These were calculated from a triple correlation using lags up to 14 bins in time and space. These values show increased contributions of motif classes with synchronous elements, as expected. C) The periodic signal is embedded in more noise (SNR = *−*9 dB), albeit still visible from the raster. D) Same as panel B with lower magnitude signals. E) The synchronous structure is embedded in more noise (SNR = *−*17 dB), but now not overtly apparent to the naked eye. F) Same as panels B and D with lower magnitude signals. All motif-class contributions are closer to 0, but motif classes with synchronous elements are still detected. G) The synchronous structure is now embedded in extreme noise (SNR = *−*40 dB). H) Same as panels B, D, and F, but now other motif-class signals are also detected (and not just those with synchronous elements) due to chance. All motif classes have even lower magnitude signals, approaching pure noise.

### 3.3 Detecting various spiking sequences

Next we tested that our method correctly detects various isolated spike sequences (triplets). Our approach correctly detected third-order motifs by the appropriate motif class. We illustrate six simulations, each including only one repeated triplet across the raster. In addition to motif class 0, all motif classes are composed of one or more constituent motif classes (I, III, and/or V). Of the simulations depicted, the first three show triplets whose motif classes are purely composed of a single constituent motif class (Fig. 4A-C), and the other three show motif classes composed of a mix of two different constituent motifs (Fig. 4D-F). Fig. 4A-C show that the second- and third-order motifs for local dynamics (Fig. 4A; I & II), synchrony (Fig. 4B; III & IV), and feedforward (Fig. 4C; V & XIII) are correctly detected. Note that motif class V (simple second-order cross-correlation, which we describe as “spike propagation”) is in fact a constituent part of every third-order motif other than those in motif classes II and IV, and so was detected at lower levels in every subsequent test. Fig. 4D-F shows the other three cases: feedback (IX; composed of I and V), divergence (XI; composed of III and V), and convergence (XII; composed of III and V).

**Fig. 4.**
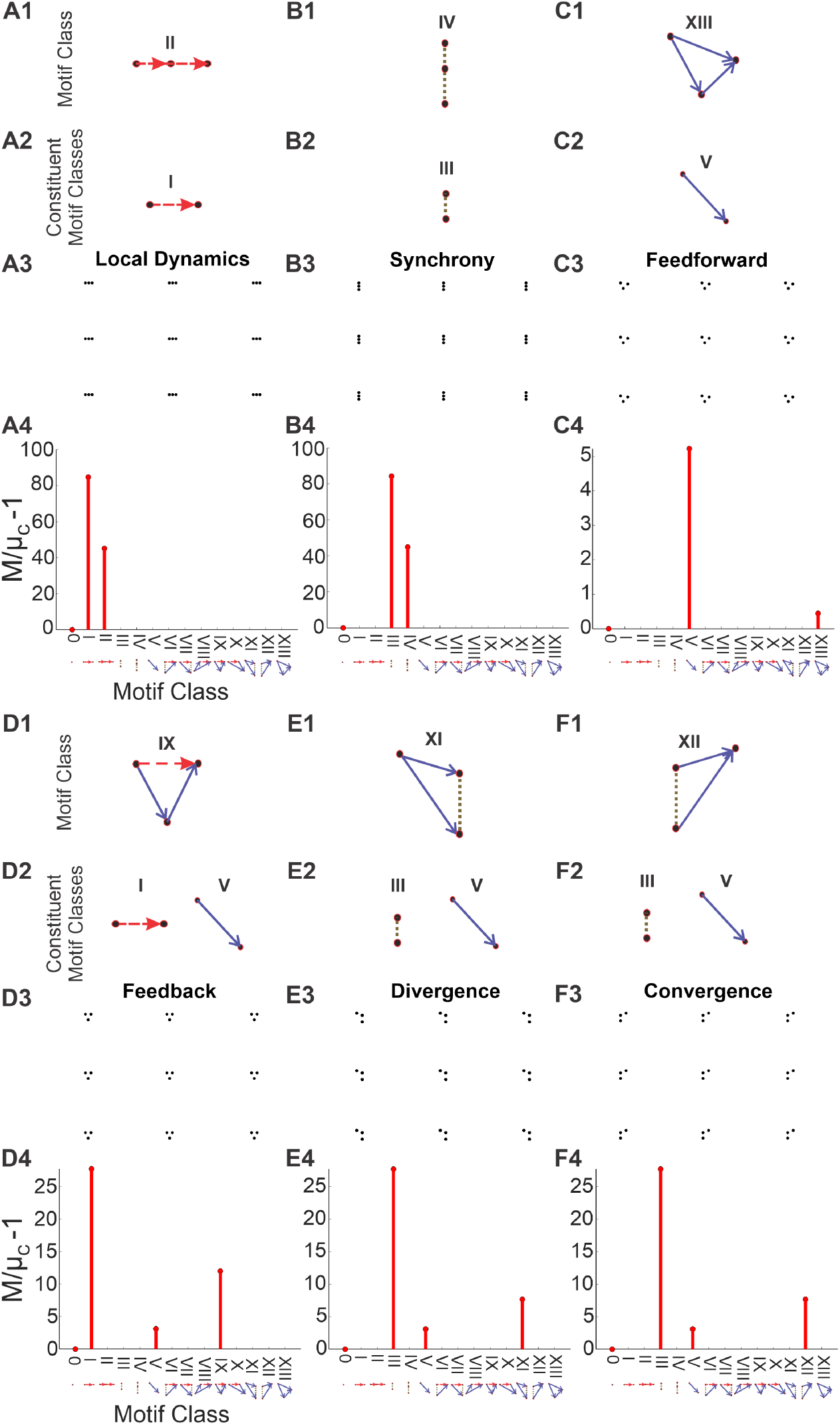
Detecting motif classes in individual patterns. We simulated 150*×*150 rasters, each consisting of a single repeated triplet. *Row 1* : the motif class of the repeated triplet. *Row 2* : the motif class’ constituent motif classes. *Row 3* : the simulated raster. *Row 4* : the motif-class contributions (*M/μ*_*c*_ − 1) calculated from the row 3 raster’s triple correlation, using lags of up to 14 bins (which is less than the separation between motifs in the raster). In all cases, the highest-order motif with a non-zero contribution is the motif class, and the remaining non-zero contributions are the constituent motif classes. Note the changing y-axis scale: motif classes I and III are far less likely to occur from chance than motif class V.

In every case, including those not depicted, the tested triplets were reflected by the correct motif-class contribution (along with the triplets’ constituent parts). For example, a feedback triplet elevates the contribution from motif-class IX, along with contributions from motif-classes I and V, which together constitute motif-class IX (Fig. 4D). Note that the particular summary used in this approach does not depend on quantitative details. Because of this, our detector works purely on 1) the qualitative spike sequencing that defines the motif-class, and 2) the presence of that motif class anywhere within the raster.

### 3.4 Application to experimental data

We further tested our approach using open-source, publicly available network data recorded from rat cortical cultures (details of this dataset are described in [20] and obtained from [21]). Briefly, rat cortical neurons were cultured in microelectrode array (MEA) well plates consisting of 16 (4×4 array) electrodes.

Data were collected at 22 days *in vitro*. The datasets consisted of already-detected spike time data from baseline wells (*n* = 35 wells) and from wells treated with the following pharmacological agents (*n* = 35 wells):

1. Control (n=7 wells)
2. *γ*-aminobutyric acid (GABA, 10 μM, n=7 wells)
3. 6-cyano-7-nitroquinoxaline-2,3-dione (CNQX, 50 μM, n=7 wells), an *α*-amino-3-hydroxy-5-methyl-4-isoxazolepropionic acid (AMPA)/kainate receptor antagonist
4. D-(-)-2-amino-5-phosphonopentanoic acid (D-AP5, 50 μM, n=7 wells), an N-methyl-D-aspartate (NMDA) receptor antagonist
5. Gabazine (30 μM, n=7 wells), a GABA_A_ receptor antagonist

The raw data were sampled at 12,500 samples/s/channel, and the spikes were detected by the researchers as described in [21]. We downsampled the spike rasters to 500 samples/s/channel. Where multiple spikes were occasionally binned together, we counted them as a single spike. Representative 1-minute snippets of each raster per condition are depicted in Fig. S1. We computed the triple correlation for each well of the MEA plate with temporal lags of −50ms:50ms and spatial lags of −1:2 electrodes^2^ in each of the two dimensions of the 4×4 array. We first show that the results from the triple correlation approach to quantify spike rate (contributions of motif class 0, Fig. 5A) concur with the results reported in [20] (their Fig. 7C). Then for each pair of baseline and treatment wells, we calculated the ratio between *M/μ*_*c*_ – 1 for the treatment well over the baseline well, and reported these values with a boxplot for each treatment (Fig. 5B-D).

**Fig. 5.**
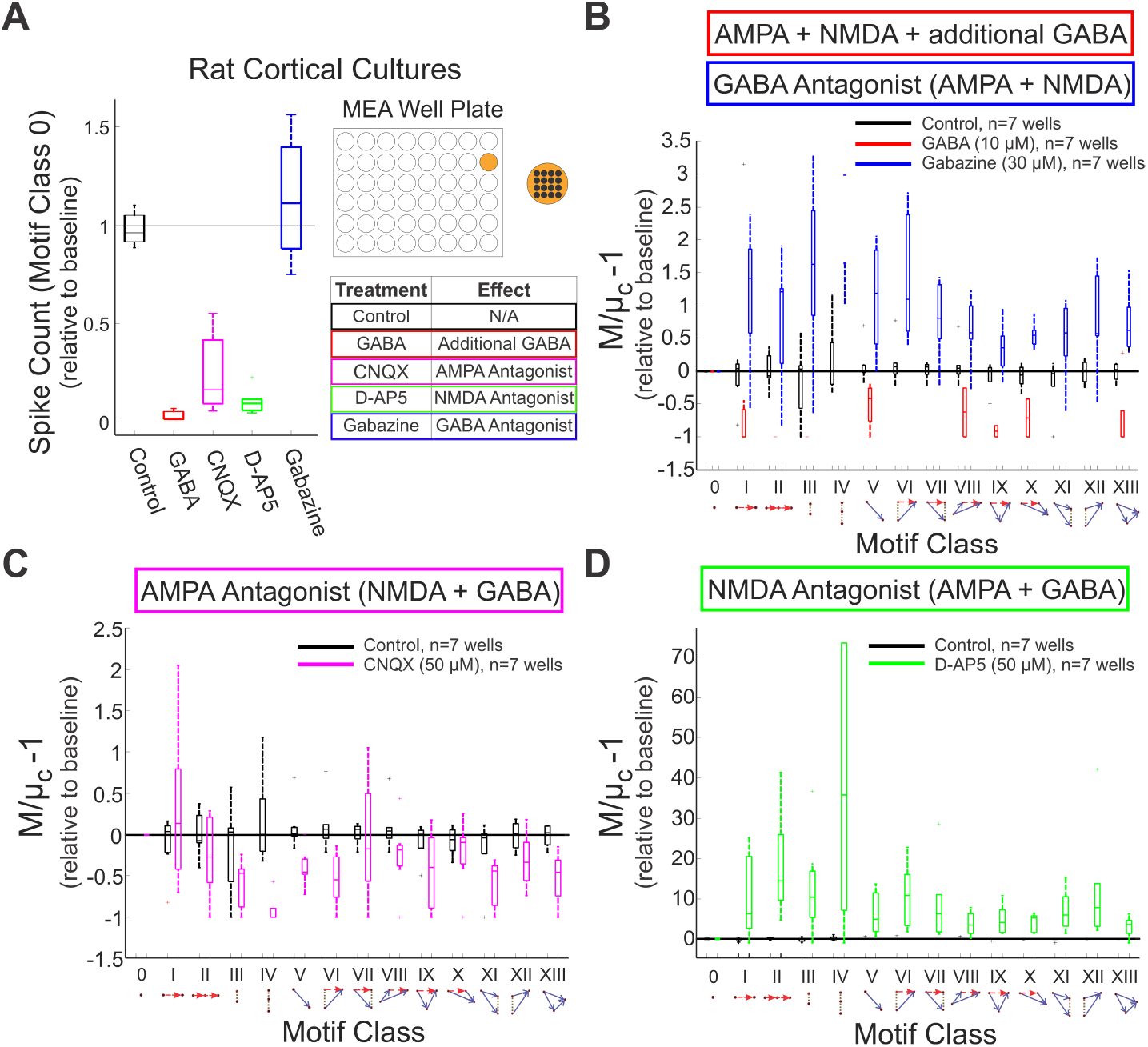
Application of the triple correlation approach to experimental data. We determined motif-class contributions in an open-source dataset of rat cortical cultures [21]. Networks (*n* = 70 wells; *n* = 35 baseline and *n* = 35 treated) were cultured on microelectrode array (MEA) well plates—each well configured with 4*x*4 electrodes. The treated wells were exposed to the following experimental conditions (*n* = 7 wells per condition): control (black), GABA (red), gabazine (blue), CNQX (magenta), and D-AP5 (green). Each treatment well was matched to an untreated baseline well. For each motif class, we depict the distribution of ratios between the treatment and baseline wells. A) The normalized spike counts relative to baseline (contributions of motif class 0) are shown for each experimental condition. The table shows the effect of each of the experimental conditions. B) *M/μ*_*c*_ *−* 1 values are shown for control-, GABA-, and gabazine-treated cultures. Note that some motif classes (IV, VI, VII, XI, XII) do not show values for GABA treatment due to lack of spiking. C) Results for control- and CNQX-treated cultures. D) The motif-class spectra for control and D-AP5-treated cultures indicate increased network structure for the latter, while its level of activity was reduced (Panel A).

The drugs used in this study provide specific levers to control synaptic activity: GABA increases and gabazine decreases inhibitory synaptic function; CNQX blocks faster excitatory synapses (AMPA/kainate); and D-AP5 blocks slower excitatory synapses (NMDA). These have broadly straightforward effects on the network’s firing rate: potentiating inhibitory synapses or blocking excitatory synapses decreases the firing rate, while blocking inhibitory synapses increases the firing rate (Fig. 5A; [20]) Modulating the inhibitory synapses has a similar gross effect: excess inhibition suppresses contributions throughout, while decreasing inhibition leads to an increase in structured firing across the motif-class spectrum (Fig. 5B). When antagonizing particular excitatory receptors—with either faster or slower post-synaptic potentials— the triple correlation reflects changes in network behavior beyond simple firing rate modulation. On top of the gross reduction in firing, antagonizing faster excitatory AMPA/kainate receptors results in a decrease in synchrony and (most) higher-order motifs involving synchrony (Fig. 5C). On the other hand, antagonizing slower excitatory NMDA synapses, creating a network in which excitation is governed by faster synapses, results in a marked increase across the motif-class spectrum, despite the overall reduction in activity. The particularly high prevalence of third-order synchrony (motif class IV) reflects the fact that the remaining firing is overwhelmingly synchronous, well above the expectation governed by chance (Figs. 5D, S1). These results agree with the connectivity graphs comparing faster and slower correlations in AMPA and NMDA networks ([22], their Fig. 9).

## 4 Discussion

The essence of this study is the introduction of a new analytical tool that fully characterizes network activity using triplet motifs. Our mathematically unique statistical analysis not only encompasses typical first- and second-order analyses (e.g. auto-correlation, cross-correlation; Fig. 1), but also includes third-order correlations that are reflective of nonlinear network behaviors. In the preceding section, we demonstrated the robustness of our approach with networks of escalating complexity: in a simple synchronous test case (Fig. 2), in the face of substantial noise (Fig. 3), and in an experimental dataset (Fig. 5). We also validated our simple summary metric in response to all computationally relevant motif classes (Fig. 4). The beauty of this approach lies in its flexible application to a multitude of finite data sets, including spike raster plots, fMRI images, local field potential (LFP), electroencephalogram (EEG), and data analyses in many other disciplines. Thus, we are excited to introduce this new analytical approach and eagerly anticipate its use as a tool to uncover new insights into network behavior.

Historically, neural network activity has been an important topic of investigation [23]. Hebb’s foundational idea of cell assemblies, which aimed to link physiology and function [24], have often been investigated by searching for repeated patterns of spiking. Typically, these cell assemblies are investigated in the context of searching for multispike pattern activation that may not be time-locked to any external stimulus or action, as “would be the case for internal processes like recalling a memory or planning a movement” [1]. The field has expanded considerably over the decades, fostering research in precise zero-phase lag synchronization [25, 26, 27], temporally-coded sequential patterns [28, 8], and so-called “synfire” chain-like structures [29, 30, 31, 32]. Our approach builds on this active area of research by integrating across multiple patterns within spatiotemporal motifs. Much as multispike patterns are putatively reflective of cell assemblies, these spatiotemporal motifs are potentially reflective of underlying structure, and thus, changes in structure could be indicative of network state shift, for example: the transition between normal and seizure states [33], preparation-to-movement in monkeys [34], sleep/wake states [35], or neuromodulation in response to pharmacological agents [36].

When applying our approach to investigate a particular hypothesis, there are two critical questions that should be asked: 1) what choice of spatiotemporal lags is relevant for the dataset (e.g. are there *a priori* synaptically relevant temporal lags?); and 2) depending on the null hypothesis, what choice of method for random chance firing is best suited for the dataset? The former presents an exciting and unexplored frontier that allows researchers to tailor this approach to a variety of hypotheses. The latter can be informed by prior literature either on null hypothesis distributions (to calculate theoretical expectations) or surrogate generation (to estimate the same). In order to present a foundational concept, we used Poisson processes to model our neurons, both in theory and in surrogate datasets. For future applications, when testing specific hypotheses using experimental data, a more nuanced surrogate (e.g. [37, 38, 39]) should be used: e.g. a stimulated neuron will not obey a simple Poisson process. Thus, the particular surrogate would depend on the hypothesis being tested and must match the null hypothesis.

Among signal processing tools, a prominent example of another unique transform of neural data is the Fourier transform, with amplitudes and phases of frequencies as fundamental units [40]. Much of medical imaging relies on the uniqueness of the Fourier transform, including computerized tomography (CT) and magnetic resonance imaging (MRI), which uses the Fourier transform to generate images [40]. In EEG and LFP research, application of Fourier transform has progressed our understanding based in the frequency spectrum [40]. Since triple correlation also constitutes a unique characterization of neural data, we present our approach as a more complex yet still useful tool, as it is not only nonlinear and higher-order, but also comprises fundamental units, triplet motifs, that are still intuitively informative. Analogous to the typical summary of the Fourier transform’s frequency spectrum into frequency bands (alpha, beta, gamma, etc.), we summarize the triple correlation’s motifs into a spectrum of fourteen motif classes. However, unlike the EEG frequency bands, which depend on clinically defined ranges, the motif-class spectrum arises directly from theory and is purely derived from possible spike sequences. It is a natural summary, and the classes themselves reflect their parsimony: they distinguish fundamental properties of computation, such as synchrony (motif classes III and IV), feedback (motif class IX), etc. (Fig. 1C). Furthermore, the constituent motif classes (I, III, and V) capture the well-known second-order correlations in analyzing neural spike data. Thus, while there are many alternative summaries, ours is both natural and useful in quantifying activity patterns (as exemplified in Figs. 2-5). Curiously, these theoretically defined fundamental units agree with previously data-driven work that has pointed to the primacy of third-order network motifs [41, 2, 3, 4].

Not only is triple correlation unique in the time domain, but its own Fourier transform, the bispectrum, is unique in the frequency domain. For the spike raster, the spectral aspect also includes the well-known power spectrum and cross-spectrum (and coherence), the Fourier transform of autocorrelation (I) and cross-correlation (V) respectively (e.g. [40]). The bispectrum’s uniqueness gives theoretical weight to the growing consensus among researchers that insights about neural signals are encoded in the bispectrum [42, 43]. Specifically, this underscores the importance of the fundamental elements of the bispectrum, which are the relationships between frequencies’ amplitudes and phases. In some cases, as in our simulated rasters (Figs. 2-4), the spatial frequencies may not be relevant due to the arbitrary ordering of neuron rows in the rasters. In other cases, where the spatial dimension has a real ordering (e.g. our MEA dataset; Fig. 5) the spatial frequency bands can be an untapped source of insight.

Both triplet motifs and inter-frequency analyses are already important topics of investigation in neuroscience (e.g. [6, 7, 41, 2, 5, 44]). Here, we have provided a direct avenue for these investigations and proven the fundamental importance of these topics to any spatiotemporal neural data. Furthermore, the success of using the bispectrum as an input to artificial neural network seizure classifiers [42] suggests that the equivalently-unique and also-meaningful triple correlation might also prove a good feature space on which to train machine learning algorithms. While we applied triple correlation to simulated and experimental spiking activity, our methodology can extend beyond spike rasters to even higher dimensionality rasters and to continuous-valued signals, such as multi-electrode LFP data or EEG recordings (see proof section “Proof in Online Materials”). Ultimately, just as frequency bands have been considered fundamental components of brain activity with Fourier transform, here we propose triplet motifs as new fundamental building blocks, one step more complex, that hold promise as an innovative approach to analyzing spatiotemporal neural activity across the breadth of recording modalities.

## Data availability

All data used is publicly available from [21].

## Code availability

The code used to generate all the results and figures in this paper is available on Github in the repository grahamas/RasterUniqueCharacterizationCode.

## Acknowledgments

We would like to thank Andrew K. Tryba, PhD, Rashi Bhatt, Benjamin Wang, Jason N. MacLean, PhD, and Stephan A. van Gils, PhD for thoughtful discussion and feedback. W.v.D. is supported by NIH Grant R01 NS-084142. S.S.D. is supported by University of Chicago MSTP Training Grant T32GM007281. G.A.S. is supported by the Pritzker Endowment for the Neurosciences.

## Footnotes

1 The theory was invoked in arguments about the Julesz conjecture [11, 12, 13, 14, 15], which stated that infinite textures with identical third-order correlations are indistinguishable to a human observer. As noted by [15], these arguments were made somewhat convoluted by the fact that the Julesz conjecture concerns textures rather than images. A mathematical texture is a statistical ensemble of images (i.e. an infinite collection of images), whereas an image is finite and bounded.

2 The spatial lags are asymmetric in order to cover the MEA without overlap, given periodic boundary conditions.

