## Supplementary Materials for "Third-order motifs are sufficient to fully and uniquely characterize spatiotemporal neural network activity"

**The PDF file includes:**

Online Methods

Tables S1 & S2

Figure S1

Reference 45

### Online Methods

#### Uniqueness of Triple-Correlation for Network Spiking Activity

We represent neuronal activity as a typical two-dimensional raster, $x(n,t)$, where $n$ is neuron location and $t$ is time. We assume that the raster is a binary (black and white, cf. Fig. 1A), meaning $x(n,t) = 1$ if neuron $n$ fires at time $t$, and $x(n,t) = 0$ otherwise. We note that the reasoning in the proofs below works just as well for any raster taking bounded values (*13*) analogous to a greyscale image, such as would be the case with local field potential recordings or the electroencephalogram. Indeed the reasoning below applies to any finite bounded dataset.

The triple correlation of $x(n,t)$ is

$$c_3(\nu_1, \tau_1, \nu_2, \tau_2) = \iint x(n,t)x(n+\nu_1, t+\tau_1)x(n+\nu_2, t+\tau_2)\, dn\, dt \tag{1}$$

for spatial lags $\nu_1, \nu_2$ and temporal lags $\tau_1, \tau_2$. This is the triple correlation function that the TCU theorem shows uniquely characterizes the spike raster.

We define $\mathcal{F}[c_3]$ as the Fourier transform of $c_3$, which means

$$\mathcal{F}[c_3](\sigma_1,\omega_1,\sigma_2,\omega_2) = \iiiint c_3(\nu_1,\tau_1,\nu_2,\tau_2)e^{-j\sigma_1\nu_1}e^{-j\omega_1\tau_1}e^{-j\sigma_2\nu_2}e^{-j\omega_2\tau_2}\, d\nu_1\, d\tau_1\, d\nu_2\, d\tau_2 \tag{2}$$

We can rewrite this integral in terms of the Fourier transform of $x$ by substituting (1) into (2) and rearranging the integration order to find

$$\mathcal{F}[c_3](\sigma_1,\omega_1,\sigma_2,\omega_2) = \iiiint x(n,t)x(n+\nu_2,t+\tau_2)\left[\iint x(n+\nu_1,t+\tau_1)e^{-j\sigma_1\nu_1}e^{-j\omega_1\tau_1}\, d\nu_1\, d\tau_1\right] e^{-j\sigma_2\nu_2}e^{-j\omega_2\tau_2}\, d\nu_2\, d\tau_2\, dn\, dt. \tag{3}$$

The part in between the [...] in (3) can be rewritten as $e^{j\omega_1 t}e^{j\sigma_1 n}X(\sigma_1,\omega_1)$, with $X(\sigma_1,\omega_1)$ denoting the two dimensional Fourier transform of $x(\nu_1,\tau_1)$. Similarly, the part of Equation (3) for $\nu_2$ and $\tau_2$ and their double integral can be arranged to evaluate to $e^{j\omega_2 t}e^{j\sigma_2 n}X(\sigma_2,\omega_2)$. Substitution of these results in (3) results in:

$$\mathcal{F}[c_3](\sigma_1,\omega_1,\sigma_2,\omega_2) = X(\sigma_1,\omega_1)X(\sigma_2,\omega_2)\iint x(n,t)e^{j(\omega_1+\omega_2)t}e^{j(\sigma_1+\sigma_2)n}\, dn\, dt \tag{4}$$

The double integral evaluates to $X(-\sigma_1-\sigma_2, -\omega_1-\omega_2) = X^*(\sigma_1+\sigma_2, \omega_1+\omega_2)$, where the asterisk denotes a complex conjugate. Thus we see that the Fourier transform of $c_3$ is the bispectrum:

$$\mathcal{F}[c_3(\nu_1,\tau_1,\nu_2,\tau_2)] = X(\sigma_1,\omega_1)X(\sigma_2,\omega_2)X^*(\sigma_1+\sigma_2,\omega_1+\omega_2). \tag{5}$$

This relationship between triple correlation and the bispectrum is the third-order equivalent of the Wiener-Khinchin-Einstein theorem.

Since images have a finite support and all of the above integration limits are implicitly at $(-\infty, \infty)$, we can modify the results for finite support by multiplying the spatiotemporal domain data by a two-dimensional boxcar window, $w$, limited between $(-\Sigma, \Sigma, -\Omega, \Omega)$ (or any other window with that support): $x_b = x\,w$. The frequency domain results are then characterized by the Fourier transforms convolved (denoted by $\circledast$) with the boxcar's Fourier transform ($W$) (or that of the window applied) denoted by: $X_B = X \circledast W$. By using this notation, the results in equations (1)–(5) would be adapted by adding the b and B subscripts.

Yellott and Iverson (*13*) show in a constructive proof that a finite image (in our case a spike raster of a finite size) can be uniquely reconstructed from its third-order correlation. These authors also show and discuss how this does not hold for images of infinite size. Here we do not further discuss this aspect because the size of a spike raster (or a snapshot of any modality of neural activity) is always finite.

One critically important message of this paper is that the time domain's triple correlation and the corresponding bispectrum in the frequency domain uniquely determine the firing pattern of a network. Yellott (*14*) presents this as the TCU theorem. If we apply Yellott's TCU theorem to a spike raster, we get the following.

**Theorem 1.** If $x(n, t)$ is a raster with bounded support and another raster $y(n, t)$ has the same triple correlation function as that of $x$, then $y(n, t) = x(n + a, t + b)$ for a pair of constants $a, b$.

**Theorem 2.** If $x(\mathbf{s})$ is a raster with bounded support in $N$ spatial dimensions and another raster $y(\mathbf{s})$ has the same triple correlation function as that of $x$, then $y(\mathbf{s}) = x(\mathbf{s} + \mathbf{a})$ for a vector of constants $\mathbf{a}$.

Here we present proofs in line with the proof in Yellott and Iverson's (*13*) for two- and N-dimensional spatiotemporal data.

#### Proof of Theorem 1 (two dimensions)

Given the equality of third-order correlation functions, we can use Equation (5) to find that

$$X(\sigma_1, \omega_1)\,X(\sigma_2, \omega_2)\,X(-\sigma_1 - \sigma_2, -\omega_1 - \omega_2) = \\ Y(\sigma_1, \omega_1)\,Y(\sigma_2, \omega_2)\,Y(-\sigma_1 - \sigma_2, -\omega_1 - \omega_2) \quad (6)$$

To borrow some convenient results from probability theory (see any introductory text, e.g. (*16*)), we note that $X$ and $Y$ can be considered characteristic functions since we can consider $x$ and $y$ probability distributions: as finite images, $x$ and $y$ are bounded and nonnegative, and without loss of generality we can normalize them such that their integral is $1$. Characteristic functions (the Fourier transforms of probability distributions) have two properties that are convenient for our purposes, the first of which is that they are non-zero in a region around the origin, thus allowing us the following division (for further considerations of rigor in the one dimensional case, cf. (*45*)):

$$\frac{X(\sigma_1,\omega_1)\,X(\sigma_2,\omega_2)}{Y(\sigma_1,\omega_1)\,Y(\sigma_2,\omega_2)} = \frac{Y(-\sigma_1-\sigma_2,-\omega_1-\omega_2)}{X(-\sigma_1-\sigma_2,-\omega_1-\omega_2)}. \tag{7}$$

Next we rewrite both complex functions in terms of their amplitude and phase, i.e. $X(\sigma,\omega) = |X(\sigma,\omega)|e^{j\phi(X(\sigma,\omega))}$, where $\phi$ gives the phase. So then we can rewrite the right-hand side of (7)

$$\frac{X(\sigma_1,\omega_1)\,X(\sigma_2,\omega_2)}{Y(\sigma_1,\omega_1)\,Y(\sigma_2,\omega_2)} = \frac{|Y(-\sigma_1-\sigma_2,-\omega_1-\omega_2)|e^{j\phi(Y(-\sigma_1-\sigma_2,-\omega_1-\omega_2))}}{|X(-\sigma_1-\sigma_2,-\omega_1-\omega_2)|e^{j\phi(X(-\sigma_1-\sigma_2,-\omega_1-\omega_2))}}. \tag{8}$$

Here we use the second convenient property of characteristic functions, namely that they are Hermitian, i.e. $X(-\sigma,-\omega) = X^*(\sigma,\omega)$. By setting $\sigma_2 = 0, \omega_2 = 0$ in (8), we can use this Hermitian property to derive the fact that $|X(\sigma_1,\omega_1)|^2 = |Y(\sigma_1,\omega_1)|^2$ for any $\sigma_1,\omega_1$. In particular, $|X(-\sigma_1-\sigma_2,-\omega_1-\omega_2)| = |Y(-\sigma_1-\sigma_2,-\omega_1-\omega_2)|$ so we can flip those terms.

$$= \frac{|X(-\sigma_1-\sigma_2,-\omega_1-\omega_2)|e^{j\phi(Y(-\sigma_1-\sigma_2,-\omega_1-\omega_2))}}{|Y(-\sigma_1-\sigma_2,-\omega_1-\omega_2)|e^{j\phi(X(-\sigma_1-\sigma_2,-\omega_1-\omega_2))}} \tag{9}$$

We can also rewrite those same terms thanks to the same Hermitian property.

$$= \frac{|X^*(\sigma_1+\sigma_2,\omega_1+\omega_2)|e^{j\phi(Y^*(\sigma_1+\sigma_2,\omega_1+\omega_2))}}{|Y^*(\sigma_1+\sigma_2,\omega_1+\omega_2)|e^{j\phi(X^*(\sigma_1+\sigma_2,\omega_1+\omega_2))}} \tag{10}$$

Simple complex properties are $|X| = |X^*|$ and $\phi(X) = -\phi(X^*)$, which give us

$$= \frac{|X(\sigma_1+\sigma_2,\omega_1+\omega_2)|e^{j\phi(X(\sigma_1+\sigma_2,\omega_1+\omega_2))}}{|Y(\sigma_1+\sigma_2,\omega_1+\omega_2)|e^{j\phi(Y(\sigma_1+\sigma_2,\omega_1+\omega_2))}} \tag{11}$$

Next, we return from the amplitude and phase notation to the complex functions themselves:

$$= \frac{X(\sigma_1+\sigma_2,\omega_1+\omega_2)}{Y(\sigma_1+\sigma_2,\omega_1+\omega_2)}. \tag{12}$$

Now we define $H(\sigma,\omega) = \frac{X(\sigma,\omega)}{Y(\sigma,\omega)}$ so we can rewrite (12) as

$$H(\sigma_1,\omega_1)H(\sigma_2,\omega_2) = H(\sigma_1+\sigma_2,\omega_1+\omega_2). \tag{13}$$

Therefore by a basic result of complex analysis $H(\sigma,\omega) = e^{j(a\sigma+b\omega)}$, and thus

$$Y(\sigma,\omega) = X(\sigma,\omega)e^{j(a\sigma+b\omega)}. \tag{14}$$

This holds in a region near the origin, and another property of characteristic functions is that, if their probability distribution has finite support, the probability distribution is uniquely determined by the value of the characteristic function in a region of the origin. Thus, in the spatiotemporal domain,

$$y(n,t) = x(n+a,t+b) \text{ with constants } a, b. \tag{15}$$

#### Proof of Theorem 2 ($N$ dimensions)

Here we extend the preceding proof to $N$ dimensions. This would be necessary for analysing many clinical and experimental data sources, e.g. the plane of a two-dimensional micro-electrode array produces $2+1$ dimensional data (as in our example presented in Fig. 5). Even higher dimensions may be useful in the case where long-range connections can be represented as a hidden dimension (e.g. orientation tuning in V1). The proof is generic enough that in practice the result applies to all finite datasets.

Note that the number of motifs (i.e. the size of the triple correlation matrix) scales with exponent $2N+2$.

We define the $N+1$-dimensional spatiotemporal vector variable $\mathbf{s} = (x_1, x_2, \ldots, x_N, t)$. Then we notate the triple correlation of an $N$ dimensional raster as $c_3(\mathbf{s}_1, \mathbf{s}_2)$. We notate the $N+1$-dimensional variable's Fourier transform as

$$\mathcal{F}[\mathbf{s}] = \mathbf{\Omega} = (\sigma_1, \sigma_2, \ldots, \sigma_N, \omega), \tag{16}$$

and the corresponding bispectrum as $c_3(\mathbf{\Omega}_1, \mathbf{\Omega}_2)$.

Since the properties of characteristic functions that are key to this proof still hold in $N$ dimensions (see any introductory probability theory text, e.g. (*16*)), the proof proceeds identically to that in two dimensions, but with vector notation.

Given the equality of third-order correlation functions, we can use Equation (5) to find that

$$X(\mathbf{\Omega}_1)\,X(\mathbf{\Omega}_2)\,X(-\mathbf{\Omega}_1-\mathbf{\Omega}_2) = Y(\mathbf{\Omega}_1)\,Y(\mathbf{\Omega}_2)\,Y(-\mathbf{\Omega}_1-\mathbf{\Omega}_2) \tag{17}$$

To borrow some convenient results from probability theory, we note that $X$ and $Y$ can be considered characteristic functions since we can consider $x$ and $y$ probability distributions: as finite images, $x$ and $y$ are bounded and nonnegative, and without loss of generality we can normalize them such that their integral is $1$. Characteristic functions (the Fourier transforms of probability distributions) have two properties that are convenient for our purposes, the first of which is that they are non-zero in a region around the origin, thus allowing us the following division (for further considerations of rigor in the one dimensional case, cf. (*45*)):

$$\frac{X(\mathbf{\Omega}_1)\,X(\mathbf{\Omega}_2)}{Y(\mathbf{\Omega}_1)\,Y(\mathbf{\Omega}_2)} = \frac{Y(-\mathbf{\Omega}_1-\mathbf{\Omega}_2)}{X(-\mathbf{\Omega}_1-\mathbf{\Omega}_2)}. \tag{18}$$

Next we rewrite both complex functions in terms of their amplitude and phase, i.e. $X(\mathbf{\Omega}) = |X(\mathbf{\Omega})|e^{j\phi(X(\mathbf{\Omega}))}$, where $\phi$ gives the phase. So then we can rewrite the right-hand side of (18)

$$\frac{X(\mathbf{\Omega}_1)\,X(\mathbf{\Omega}_2)}{Y(\mathbf{\Omega}_1)\,Y(\mathbf{\Omega}_2)} = \frac{|Y(-\mathbf{\Omega}_1-\mathbf{\Omega}_2)|e^{j\phi(Y(-\mathbf{\Omega}_1-\mathbf{\Omega}_2))}}{|X(-\mathbf{\Omega}_1-\mathbf{\Omega}_2)|e^{j\phi(X(-\mathbf{\Omega}_1-\mathbf{\Omega}_2))}}. \tag{19}$$

Here we use the second convenient property of characteristic functions, namely that they are Hermitian, i.e. $X(-\mathbf{\Omega}) = X^*(\mathbf{\Omega})$. By setting $\mathbf{\Omega}_2 = \mathbf{0}$ in (19), we can use this Hermitian property to derive the fact that $|X(\mathbf{\Omega}_1)|^2 = |Y(\mathbf{\Omega}_1)|^2$ for any $\mathbf{\Omega}_1$. In particular, $|X(-\mathbf{\Omega}_1 - \mathbf{\Omega}_2)| = |Y(-\mathbf{\Omega}_1 - \mathbf{\Omega}_2)|$ so we can flip those terms.

$$= \frac{|X(-\mathbf{\Omega}_1 - \mathbf{\Omega}_2)|e^{j\phi(Y(-\mathbf{\Omega}_1 - \mathbf{\Omega}_2))}}{|Y(-\mathbf{\Omega}_1 - \mathbf{\Omega}_2)|e^{j\phi(X(-\mathbf{\Omega}_1 - \mathbf{\Omega}_2))}} \tag{20}$$

We can also rewrite those same terms thanks to the same Hermitian property.

$$= \frac{|X^*(\mathbf{\Omega}_1 + \mathbf{\Omega}_2)|e^{j\phi(Y^*(\mathbf{\Omega}_1 + \mathbf{\Omega}_2))}}{|Y^*(\mathbf{\Omega}_1 + \mathbf{\Omega}_2)|e^{j\phi(X^*(\mathbf{\Omega}_1 + \mathbf{\Omega}_2))}} \tag{21}$$

Simple complex properties are $|X| = |X^*|$ and $\phi(X) = -\phi(X^*)$, which give us

$$= \frac{|X(\mathbf{\Omega}_1 + \mathbf{\Omega}_2)|e^{j\phi(X(\mathbf{\Omega}_1 + \mathbf{\Omega}_2))}}{|Y(\mathbf{\Omega}_1 + \mathbf{\Omega}_2)|e^{j\phi(Y(\mathbf{\Omega}_1 + \mathbf{\Omega}_2))}} \tag{22}$$

Next, we return from the amplitude and phase notation to the complex functions themselves:

$$= \frac{X(\mathbf{\Omega}_1 + \mathbf{\Omega}_2)}{Y(\mathbf{\Omega}_1 + \mathbf{\Omega}_2)}. \tag{23}$$

Now we define $H(\mathbf{\Omega}) = \frac{X(\mathbf{\Omega})}{Y(\mathbf{\Omega})}$ so we can rewrite (23) as

$$H(\mathbf{\Omega}_1)H(\mathbf{\Omega}_2) = H(\mathbf{\Omega}_1 + \mathbf{\Omega}_2). \tag{24}$$

Therefore by a basic result of complex analysis $H(\mathbf{\Omega}) = e^{j\mathbf{k}\cdot\mathbf{\Omega}}$, and thus

$$Y(\mathbf{\Omega}) = X(\mathbf{\Omega})e^{j\mathbf{k}\cdot\mathbf{\Omega}}. \tag{25}$$

This holds in a region near the origin, and another property of characteristic functions is that, if their probability distribution has finite support, the probability distribution is uniquely determined by the value of the characteristic function in a region of the origin. Thus, in the spatiotemporal domain,

$$y(\mathbf{s}) = x(\mathbf{s} + \mathbf{a}) \text{ with vector of constants } \mathbf{a}. \tag{26}$$

#### Computing the triple correlation

For computational applications presented here, we consider third-order correlation analysis applied to a discrete raster with neuron rows $n$, `1:N`, and time columns $t$, `1:T`, where each pixel $r(n,t)$ is filled with a $0$ (white for no activity) or $1$ (black for spike). The complete triple correlation function, while formally requisite, incurs substantial computational cost while adding increasingly noisy information. We limit our computation to lag windows $-W_t : W_t$ and $-W_n : W_n$ chosen such that two local synaptic connections could not take longer than $W_t$, and two neurons are unlikely to be connected further than $W_n$. With this restricted lag window, we do not zero pad our raster (as in (*13*)) but instead use boundary conditions periodic in space (since our spatial ordering was already arbitrary[1]) and restrict our calculation to a subset of time such that no motifs extend beyond time zero or the raster's duration. In this case, discrete equivalent $\mathtt{cd}_3$ of the triple correlation expression $c_3$ (Equation 1) or the short form reported in the main text, $c_3(n_1, t_1, n_2, t_2) = \langle r(n,t) r(n+n_1, t+t_1) r(n+n_2, t+t_2)\rangle_{n,t}$, is

$$\mathtt{cd}_3(n_1, t_1, n_2, t_2) = \tag{27}$$

$$\frac{1}{\#(W_n, W_t, N, T)} \sum_{t=1+W_t}^{T-W_t} \sum_{n=1}^{N} \tag{28}$$

$$r(n,t) r((n+n_1-1)\%N+1, t+t_1) r((n+n_2-1)\%N+1, t+t_2).$$

We scale by the number of spike bins in the summation

$$\#(W_n, W_t, N, T) = (T - 2W_t)(N). \tag{29}$$

Equations (27) and (29) were used in all our simulations.

Note that in the text we report triple correlations "using lags up to $X$ in time and space." This corresponds to $W_n = X/2$ and $W_t = X/2$. When $X$ is odd, we use the more general lag windows $-\mathtt{floor}(X/2) : \mathtt{ceil}(X/2)$, with corresponding changes to the summation bounds and scaling instead by $(T-X)(N)$. We report the maximum lag as $X$ because, e.g., $\mathtt{cd}_3(-10, t_1, 10, t_2)$ has a maximum temporal separation between spikes of 20, despite the fact that technically the lag arguments are at most 10.

#### Summarizing triple correlation into motif classes

We summarized the triple correlation as fourteen motif-classes (Fig. 1). The main text discusses the reasoning for this particular choice, and here we describe the details.

Let $[n_1, t_1 | n_2, t_2]$ denote a motif, i.e. an argument to triple correlation, which would have a value $c_3(n_1, t_1, n_2, t_2)$. We want to group together these motifs according to what we can qualitatively distinguish. We only hold two qualitative distinctions:

[1]In the MEA case, we also use periodic spatial boundary conditions even though the ordering is not arbitrary. Zeropadding introduces problems as spikes near the center of the array are more "valuable" than spikes near the boundaries (they contribute to more motifs). An alternative would be to introduce noise past the boundaries, but that creates its own problems in the choice of noise. Since our analyses already assumed a lack of spatial information (in our choice of motif class) we decided to simply discard the spatial information for the purposes of these analyses, and therefore to use the spatial periodic boundary conditions.

1. temporal: we distinguish between before, simultaneous, and after (temporal coordinate less than, equal to, or greater)
2. spatial: we distinguish between the same and different (spatial coordinate equal and spatial coordinate not equal)

We will interpret the motif as a three node graph: one base node implicitly (0,0), and two other nodes, $(n_1, t_1)$ and $(n_2, t_2)$. Note that neither of our qualitative distinctions involve node identity. So, for example, we would not distinguish between $[0, 1|0, 2]$ and $[0, 2|0, 1]$, even though one the first node follows the second, and in the other the reverse is true. In both, the motif includes nodes at the same spatial coordinate in sequence, so we do not distinguish between them.

By grouping motifs that are indistinguishable under these criteria, we develop motif classes. We enumerate these motif classes in Figure 1. From that enumeration, it is clear that no motif can belong to two motif classes. To show that this enumeration is complete (there are no more motif classes and all motifs fall into one of these classes), we count the number of motif classes below. To begin, we count "lag-sign" motifs, which is to say, if we only care about the sign of the lags, how many motifs are there. This first step avoids some of the small complications in our definition, i.e. the different treatments of space and time, as well as the subtle ways they interact when we discard node identity. Having counted lag-sign motifs, we then reduce these lag-sign motifs by discounting spatial direction and node identity to finally arrive at the number of motif classes.

#### The number of lag-sign motifs

Given that three node combinations determine $c_3$, we want to know how many ways there are to order them in time and space, with the caveat that we may also place one node on top of another in either time, or space, or both (to account for zero lag). To count this, we divide the cases according to how many different temporal and spatial bins the nodes have between them. For example, if all the nodes are completely distinct in time and space, then there are both three spatial bins and three temporal bins between the three nodes. On the other hand, if we overlap all three nodes in both time and space, then there is only one spatial bin and one temporal bin between them. All the cases are labeled in Table S1. Note that the cases are symmetric, so we will calculate the values as if in the lower left half of the table, i.e. where the number of spatial bins is greater than or equal to the number of time bins. The arrangement of this table is the same as in Figure 1C. As described above, we call this arrangement of nodes separated by lags a motif. When we only note the sign of the lags, we call it a lag-sign motif.

A. In the case where no nodes share a spatial bin (neuron), there are $3!$ ways to order them in space. Similarly if the nodes share no time bins, then there are $3!$ ways to order the time bins. So, if no spatial bins nor time bins are shared, then there are $3!3! = 36$ orderings.

B. Given that the nodes only have two distinct spatial bins between them, there are three ways to choose which nodes share the same spatial bin, and then two ways to choose whether the remaining node has a spatial bin before or after the identical nodes' spatial bin, for $3 \cdot 2 = 6$ spatial orderings. As above, if no time bins are identical, then there are 3! temporal orderings. So there are $3 \cdot 2 \cdot 3! = 36$ orderings.

C. If nodes share the same spatial bin, then there is only one way to order that neuron. As above, if no time bins are identical then there are 3! temporal orderings, giving a total of $1 \cdot 3! = 6$ orderings. The same is true in the symmetric case when the nodes share the same time bin, but all have different neurons.

D. We already established that for the case where two spatial bins are the same, there are $3 \cdot 2 = 6$ orderings. The same is true when the nodes only have two distinct time bins. So there are $3 \cdot 2 \cdot 3 \cdot 2 = 36$ orderings.

E. We already established that for the case where two spatial bins are the same, there are $3 \cdot 2 = 6$ orderings, and for the case where all time bins are the same, there is only one ordering. So there are $3 \cdot 2 \cdot 1 = 6$ orderings.

F. In the case where all spatial bins and all time bins are the same, there is only one ordering.

We sum up the table: $A + 2B + 2C + D + 2E + F = 36 + 2 \cdot 36 + 2 \cdot 6 + 36 + 2 \cdot 6 + 1 = 169$ lag-sign motifs.

#### The number of motif classes

To complete our summary, we reduce the 169 lag-sign motifs listed in Table S2 by appealing to two symmetries: one in space, and one in permutation. The spatial one is straightforward: since we have not assigned meaning to space, reordering the spatial bins does not change our interpretation. The permutation symmetry is more technical: the way triple correlation is calculated, the ordering of the nodes matters. In second-order correlation (call it $\rho$), this is even symmetry: $\rho(x) = \rho(-x)$. More generally, a motif has some polygonal structure (or possibly a point or a line; treat these as straightforward special cases of the following argument). Each vertex of the polygon has a node identity: in the triple correlation case, one is the base node, one is the first node, given by the base node plus $(n_1, t_1)$, and one is the second node, given by the base node plus $(n_2, t_2)$. But we don't actually care which is the base, which is the first, and which is the second node: triple correlation is invariant under permutation of these nodes. We reduce the number of lag-sign motifs by accounting for these symmetries. Again working from Table S1:

A. There are 3! ways to permute node identity, and 3! ways to permute neuron identity, independently, giving a product of $36$ permutations. So all $36$ lag-sign motifs in category A reduce to 1 motif class (Fig. 1C, XIII).

Table S1: The number of possible orderings in space and time given two possible time lags and two possible spatial lags, which corresponds to up to three possible bins in both time and space. Note that in case of a spike raster of single unit activity, each spatial bin corresponds to one neuron. This table corresponds to Fig. 1C in the main text.

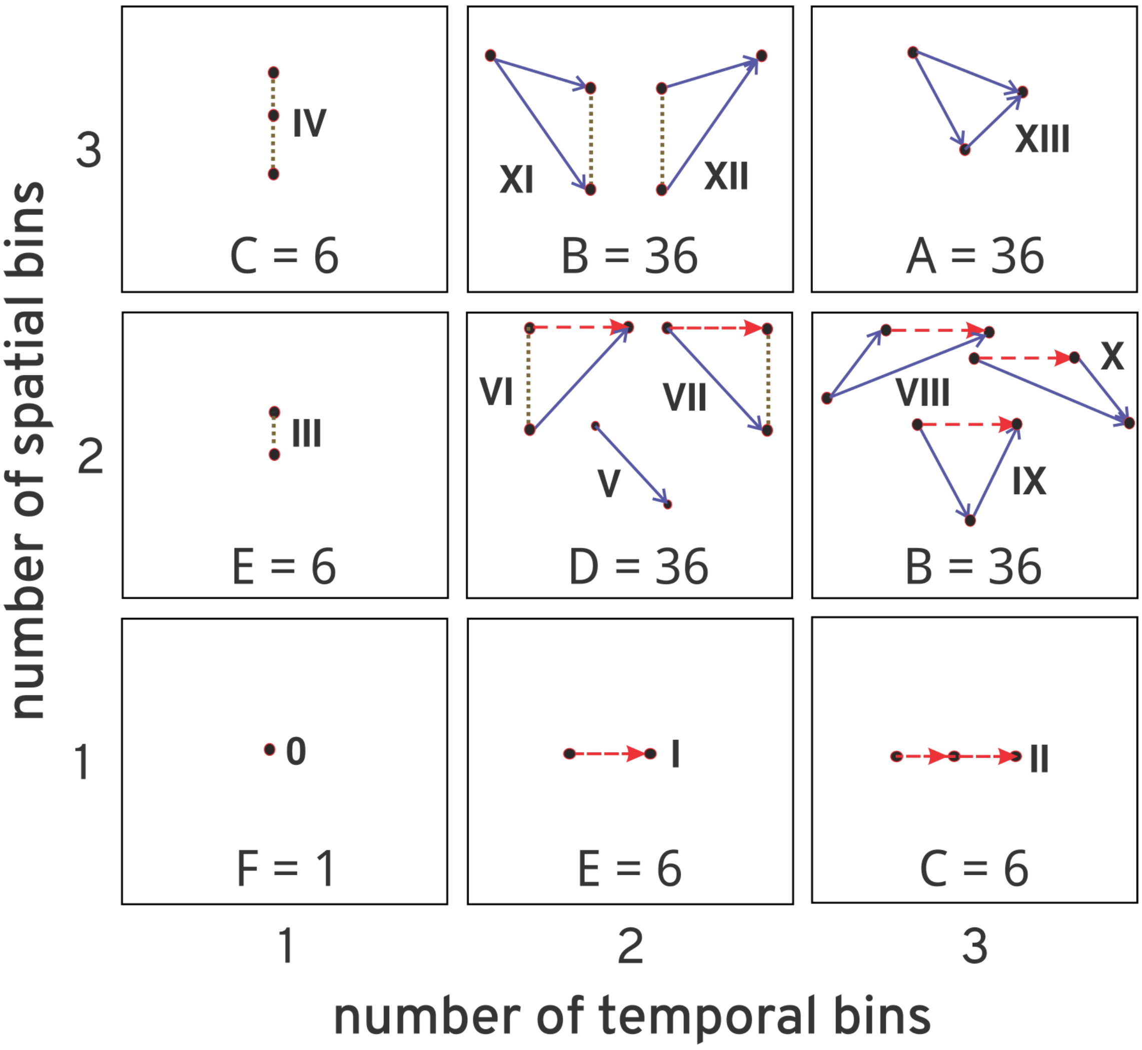

B(2T). For category B with two time bins, there are 3! node permutations, and only 3 spatial permutations: two spatial bins share a time bin, and the third spatial bin may appear above, between, or below these two spatial bins (due to the fact that we are reducing from lag-sign motifs, not from lag-motifs). The product is 18 total permutations, so the 36 lag-sign motifs in category B(2T) reduce to 2 motif classes (Fig. 1C, XI and XII).

B(2S). For category B with two spatial bins, there are only two spatial permutations: one spatial bin has two time bins, and the remaining spatial bin can either be above or below. There are still 3! node permutations, leading to a total of 12 permutations. Therefore the 36 lag-sign motifs in category B(2S) reduce to 3 motif classes (Fig. 1C, VIII, IX, and X).

D. By the same argument as in B(2S), there are two spatial permutations, and again 3! node permutations, leading to a total of 12 permutations. Therefore the 36 lag-sign motifs in category D reduce to 3 motif classes (Fig. 1C, V, VI, and VII).

C(1T). When there is only one time bin, spatial permutation is the same as node permutation, so there are only the $3! = 6$ permutations. Therefore the 6 lag-sign motifs in category C(1T) reduce to 1 motif class (Fig. 1C, IV).

E(1T). As in C(1T), all 6 reduce to 1 motif class (Fig. 1C, III).

C(1S). When there is only one spatial bin, the lag-sign motifs have no spatial permutations, only node permutations, so the 6 lag-sign motifs in category C(1S) reduce to 1 motif class (Fig. 1C, II).

E(1S). As in C(1S), all 6 reduce to 1 motif class (Fig. 1C, I)

F. In the case where all lags are zero, triple correlation has no symmetric entries but the zero entry, so there are no permutations, and the 1 lag-sign motif (which also contains only one lag-motif) remains 1 motif class (Fig. 1C, 0).

The above sum to 14 motif classes, as depicted in Figure 1.

#### Expected contributions per motif class

Because it is important to asses how these motif-class contributions be explained by chance (or not), we simulate 100 activity matched noise rasters for each raster under investigation, and compare the raster's motif-class spectrum with that of its noise matched rasters. In the following, we derive the theoretical expectations for rasters governed by noise.

Let $r$ be a raster with $N$ neurons and $T$ time bins. In a binary raster, a spike is indicated by $r(n, t) = 1$, else $r(n, t) = 0$. We calculate the triple correlation $c_3$ over the whole raster with periodic boundary conditions (i.e. $r(n+\eta, t+\tau) = r((n+\eta) \mod N, (t+\tau) \mod T)$, with an additional 1 offset). Define $\pi(\mathbf{x}, \boldsymbol{\lambda}) = r(\mathbf{x})r(\mathbf{x}+\boldsymbol{\lambda}_1)r(\mathbf{x}+\boldsymbol{\lambda}_2)$ and for shorthand $\pi(\boldsymbol{\lambda}) = \pi(0, \boldsymbol{\lambda})$. In general with lags $\boldsymbol{\lambda} = (\eta_1, \tau_1, \eta_2, \tau_2)$

$$\begin{aligned} c_3(\boldsymbol{\lambda}) &= c_3(\eta_1, \tau_1, \eta_2, \tau_2) \\ &= \sum_{(n,t)\in I(r)} r(n,t)r(n+\eta_1, t+\tau_1)r(n+\eta_2, t+\tau_2) \\ &= \sum_{\mathbf{x}\in I(r)} \pi(\mathbf{x}, \boldsymbol{\lambda}) \end{aligned}$$

where $I(r)$ is the set of all indices of $r$, in this case $I(r) = (1 : N) \times (1 : T)$. We are interested in the motif class contributions, which are the sums of triple correlations for lags that constitute each of the fourteen motifs. So, if $\boldsymbol{\lambda} \in \Delta(M_i)$ denotes that the lags $\boldsymbol{\lambda}$ constitute a triplet in motif class $i$, then we define the motif class contribution as

$$M_i(r) = \sum_{\boldsymbol{\lambda}\in\Delta(M_i)} c_3(\boldsymbol{\lambda})$$

The expectation then is given by

$$\begin{aligned} \langle M_i(r)\rangle &= \left\langle \sum_{\boldsymbol{\lambda}\in\Delta(M_i)} c_3(\boldsymbol{\lambda}) \right\rangle \\ &= \left\langle \sum_{\boldsymbol{\lambda}\in\Delta(M_i)} \sum_{x\in I(r)} \pi(x, \boldsymbol{\lambda}) \right\rangle \\ &= \sum_{\boldsymbol{\lambda}\in\Delta(M_i)} \sum_{x\in I(r)} \langle \pi(x, \boldsymbol{\lambda})\rangle \\ &= \sum_{x\in I(r)} \sum_{\boldsymbol{\lambda}\in\Delta(M_i)} \langle \pi(x, \boldsymbol{\lambda})\rangle \\ &= NT \sum_{\boldsymbol{\lambda}\in\Delta(M_i)} \langle \pi(\boldsymbol{\lambda})\rangle \end{aligned}$$

In fact, $\langle \pi(\boldsymbol{\lambda})\rangle$ is constant for all $\boldsymbol{\lambda} \in \Delta(M_i)$ for a given $i$, so

$$\langle M_i(r)\rangle = NT\#(\Delta(M_i)) \langle \pi(\lambda_i)\rangle . \tag{30}$$

We can easily calculate $\langle \pi(\boldsymbol{\lambda})\rangle$ if we assume independent bins. See below for the calculation in each of the cases of independent Bernoullis.

$\#(\Delta(M_i))$ is the more challenging problem, which we calculate case-by-case below in the section "The number of triplet motifs."

##### Independent spiking with constant probability (Bernoulli)

For a simple spiking raster, we assume all bins have independent probability $p$ of spiking, i.e. they are all independent Bernoullis.[2] This lets us substitute into equation 30 the following probabilities

$$\begin{aligned} \langle \pi(\boldsymbol{\lambda}_0) \rangle &= p \\ \langle \pi(\boldsymbol{\lambda}_i) \rangle &= p^2 \qquad \text{for } i = \mathrm{I}, \mathrm{III}, \mathrm{V} \\ \langle \pi(\boldsymbol{\lambda}_i) \rangle &= p^3 \qquad \text{for } i = \mathrm{II}, \mathrm{IV}, \mathrm{VI}, \mathrm{VII}, \mathrm{VIII}, \mathrm{IX}, \mathrm{X}, \mathrm{XI}, \mathrm{XII}, \mathrm{XIII} \end{aligned}$$

where $\boldsymbol{\lambda}_i$ is a representative triplet of motif class $i$. This representative is sufficient because the expectation of the product of bins purely depends on the number of distinct bins multiplied. Within a motif class, all triplets have the same number of distinct bins, with motif class 0 having only one spike, motif classes $\mathrm{I}, \mathrm{III}, \mathrm{and} \mathrm{IV}$ having two spikes, and the remaining motif classes having three spikes. Knowing these values, we have the expectations of each motif class.

##### The number of triplet motifs

The quantity $\#(\Delta(M_i))$ simply counts the number of motifs in a given motif class. This is independent of the data's distribution, but dependent on the data's shape. Let $\Lambda_t$ be the number of time lags considered, and let $\Lambda_t^{\pm}$, $\Lambda_t^{+}$, and $\Lambda_t^{-}$ be the number of time lags that are nonzero, positive, and negative respectively. Similarly define $\Lambda_n$, $\Lambda_n^{\pm}$, $\Lambda_n^{+}$, and $\Lambda_n^{-}$.

Our first step for each motif class will be to define $\Delta(M_i)$, for which purpose we will define a notation to indicate the sign and equality of each part of the two spatiotemporal lags (the third lag implicitly zero). For example, in the case of motif class XIII, we write $\Delta(M_{\mathrm{XIII}}) = [x^{\pm}, t^{\pm} \mid y^{\pm}, s^{\pm}]$. This indicates that all four lags are nonzero ($\pm$ superscript) and none are equal ($x \neq y$ and $t \neq s$). In contrast, we could write $[0, t^{+} \mid 0, t^{+}]$ to indicate both spatial lags are zero, and the temporal lags are both positive and equal. We add a coefficient "2" to indicate that we want to additionally include set of lags symmetric with the notated lag, i.e. flipping which lag is first and which is second, e.g. $2[0, 0 \mid 0, t^{\pm}] = [0, 0 \mid 0, t^{\pm}] + [0, t^{\pm} \mid 0, 0]$.

I Motif class I consists in lags $\Delta(M_{\mathrm{I}}) = 2[0, 0 \mid 0, t^{\pm}] + [0, t^{\pm} \mid 0, rt^{\pm}]$. We count these as follows: there is only one way to choose the $(0, 0)$ lag, and $\Lambda_t^{\pm}$ ways to choose a $(0, t^{\pm})$ lag. Therefore there are $1 \cdot \Lambda_t^{\pm}$ lags in $[0, 0 \mid 0, t^{\pm}]$. By symmetry there are the same number in $[0, t^{\pm} \mid 0, 0]$, so the coefficient 2 works as advertised, giving $2\Lambda_t^{\pm}$ lags in $2[0, t^{\pm} \mid 0, 0]$. For the second addend, there are $\Lambda_t^{\pm}$ ways to choose the $(0, t^{\pm})$ lag. Since the lags are equal there is only one choice for the

[2] This also means that all the neurons are simulated Poisson processes with identical rates.

second lag, leading to a $\Lambda_t^\pm$ lags in $[0t^\pm \mid 0t^\pm]$. In total then, there are $3\Lambda_t^\pm$ lags in $\Delta(M_I) = 2[0,0 \mid 0,t^\pm] + [0t^\pm \mid 0t^\pm]$. We write this

$$\begin{aligned}\#(\Delta(M_I)) &= \#(2[0,0 \mid 0,t^\pm] + [0t^\pm \mid 0t^\pm]) \\ &= 2(1 \cdot \Lambda_t^\pm) + (\Lambda_t^\pm \cdot 1) \\ &= 3\Lambda_t^\pm\end{aligned}$$

II For motif class II, we have $\Delta(M_{II}) = [0,t^\pm \mid 0,s^\pm]$. This time for our second lag, we have one fewer choice because we must have that the temporal lags do not equal. So we find

$$\begin{aligned}\#(\Delta(M_{II})) &= \#([0,t^\pm \mid 0,s^\pm]) \\ &= \Lambda_t^\pm(\Lambda_t^\pm - 1)\end{aligned}$$

III For motif class III, the situation is the same as motif class I, but with $n$ rather than $t$, so $\Delta(M_{III}) = 2[0,0 \mid x^\pm,0] + [x^\pm,0 \mid x^\pm,0]$, giving

$$\begin{aligned}\#(\Delta(M_{III})) &= \#(2[0,0 \mid x^\pm,0] + [x^\pm,0 \mid x^\pm,0]) \\ &= 2(1 \cdot \Lambda_n^\pm) + (\Lambda_n^\pm \cdot 1) \\ &= 3\Lambda_n^\pm\end{aligned}$$

.

IV Similarly motif class IV has $\#(\Delta(M_{III})) = \Lambda_n^\pm(\Lambda_n^\pm - 1)$.

V

$$\begin{aligned}\Delta(M_V) &= 2[0,0 \mid x^\pm,t^\pm] + [x^\pm,t^\pm \mid x^\pm,t^\pm] \\ \#(\Delta(M_V)) &= 2\Lambda_n^\pm\Lambda_t^\pm + \Lambda_n^\pm\Lambda_t^\pm \\ &= 3\Lambda_n^\pm\Lambda_t^\pm\end{aligned}$$

VI

$$\begin{aligned}\Delta(M_{VI}) &= 2[0,t^+ \mid x^\pm,0] + 2[x^\pm,0 \mid x^\pm t^+] + 2[0,t^- \mid x^\pm t^-] \\ \#(\Delta(M_{VI})) &= 2(\Lambda_t^+\Lambda_n^\pm) + 2(\Lambda_n^\pm\Lambda_t^+) = 2(\Lambda_t^-\Lambda_n^\pm) \\ &= 4(\Lambda_n^\pm\Lambda_t^+) + 2(\Lambda_n^\pm\Lambda_t^-)\end{aligned}$$

VII By symmetry with VI,

$$\#(\Delta(M_{VII})) = 4(\Lambda_n^\pm\Lambda_t^-) + 2(\Lambda_n^\pm\Lambda_t^+)$$

VIII

$$\begin{aligned}
\Delta(M_{VIII}) &= [x^{\pm}, t^{+} \mid x^{\pm}, s^{+}] + 2[x^{\pm}, t^{-} \mid 0, s^{+}] + 2[x^{\pm}, t^{-} \mid 0, s^{-}(> t^{-})] \\
\#(\Delta(M_{VIII})) &= \Lambda_n^{\pm}\Lambda_t^{+}(\Lambda_t^{+} - 1) + 2\Lambda_n^{\pm}\Lambda_t^{-}\Lambda_t^{+} + 2\Lambda_n^{\pm} \sum_{\Lambda_t^{-} < t^{-} < -1} \#s : t^{-} < s < 0 \\
&= \Lambda_n^{\pm}\Lambda_t^{+}(\Lambda_t^{+} - 1) + 2\Lambda_n^{\pm}\Lambda_t^{-}\Lambda_t^{+} + 2\Lambda_n^{\pm} \sum_{-\Lambda_t^{-} < t^{-} < -1} |t^{-}| - 1 \\
&= \Lambda_n^{\pm}\Lambda_t^{+}(\Lambda_t^{+} - 1) + 2\Lambda_n^{\pm}\Lambda_t^{-}\Lambda_t^{+} + 2\Lambda_n^{\pm} \sum_{(-\Lambda_t^{-} - 1) > t > 1} t \\
&= \Lambda_n^{\pm}\Lambda_t^{+}(\Lambda_t^{+} - 1) + 2\Lambda_n^{\pm}\Lambda_t^{-}\Lambda_t^{+} + 2\Lambda_n^{\pm}(\Lambda_t^{-} - 1)\frac{(\Lambda_t^{-} - 1) + 1}{2} \\
&= \Lambda_n^{\pm}\Lambda_t^{+}(\Lambda_t^{+} - 1) + 2\Lambda_n^{\pm}\Lambda_t^{-}\Lambda_t^{+} + \Lambda_n^{\pm}\Lambda_t^{-}(\Lambda_t^{-} - 1)
\end{aligned}$$

IX

$$\begin{aligned}
\Delta(M_{IX}) &= 2[0, t^{+} \mid x^{\pm}, s^{+} < t^{+}] + 2[0, t^{-} \mid x^{\pm}, s^{-} > t^{-}] + 2[x^{\pm}, t^{-} \mid x^{\pm}, s^{+}] \\
\#(\Delta(M_{IX})) &= 2\Lambda_n^{\pm} \sum_{1 \leqslant t \leqslant \Lambda_t^{+}} (t - 1) + 2\Lambda_n^{\pm} \sum_{-\Lambda_t^{-} \leqslant t \leqslant -1} (|t| - 1) + 2\Lambda_n^{\pm}\Lambda_t^{-}\Lambda_t^{+} \\
&= 2\Lambda_n^{\pm}\left(\Lambda_t^{+}\frac{(\Lambda_t^{+} - 1) + 0}{2}\right) + 2\Lambda_n^{\pm}\left(\Lambda_t^{-}\frac{(\Lambda_t^{-} - 1) + 0}{2}\right) + 2\Lambda_n^{\pm}\Lambda_t^{-}\Lambda_t^{+} \\
&= \Lambda_n^{\pm}\Lambda_t^{+}(\Lambda_t^{+} - 1) + \Lambda_n^{\pm}\Lambda_t^{-}(\Lambda_t^{-} - 1) + 2\Lambda_n^{\pm}\Lambda_t^{-}\Lambda_t^{+}
\end{aligned}$$

X By symmetry with VIII

$$\begin{aligned}
\Delta(M_{X}) &= [x^{\pm}, t^{-} \mid x^{\pm}, s^{-}] + 2[x^{\pm}, t^{+} \mid 0, s^{-}] + 2[x^{\pm}, t^{+} \mid 0, s^{+}(< t^{+})] \\
\#(\Delta(M_{X})) &= \Lambda_n^{\pm}\Lambda_t^{-}(\Lambda_t^{-} - 1) + 2\Lambda_n^{\pm}\Lambda_t^{+}\Lambda_t^{-} + \Lambda_n^{\pm}\Lambda_t^{+}(\Lambda_t^{+} - 1)
\end{aligned}$$

XI

$$\begin{aligned}
\Delta(M_{XI}) &= [x^{\pm}, t^{+} \mid y^{\pm}, t^{+}] + 2[x^{\pm}, 0 \mid y^{\pm}, t^{-}] \\
\#(\Delta(M_{XI})) &= \Lambda_n^{\pm}(\Lambda_n^{\pm} - 1)\Lambda_t^{+} + 2\Lambda_n^{\pm}(\Lambda_n^{\pm} - 1)\Lambda_t^{-}
\end{aligned}$$

XII

$$\begin{aligned}
\Delta(M_{XII}) &= [x^{\pm}, t^{-} \mid y^{\pm}, t^{-}] + 2[x^{\pm}, 0 \mid y^{\pm}, t^{+}] \\
\#(\Delta(M_{XII})) &= \Lambda_n^{\pm}(\Lambda_n^{\pm} - 1)\Lambda_t^{-} + 2\Lambda_n^{\pm}(\Lambda_n^{\pm} - 1)\Lambda_t^{+}
\end{aligned}$$

XIII

$$\Delta(M_{XIII}) = [x^{\pm}, t^{\pm} \mid y^{\pm}, s^{\pm}]$$
$$\#(\Delta(M_{XIII})) = \Lambda_n^{\pm}\Lambda_t^{\pm}(\Lambda_n^{\pm} - 1)(\Lambda_t^{\pm} - 1)$$

Table S2: **motif classes for triple correlation with different spatial and temporal lags.** In the triple correlation equation, there are four lags that can be either zero or non-zero. Accordingly, the rows in this table are ordered in 16 ($2^4$) groups while the subgroups within these groups, represent the arrangement of the non-zero lags. The total number of rows is 169 lag-sign motifs for three-node motifs with different lags (Table S1). Each row represents a single three-node lag-sign motif, which differentiated by the spike sequencing. The base node implicitly has lags $n_0 = 0$ and $t_0 = 0$, and the four lags of the other two nodes ($n_1$, $n_2$, $t_1$, $t_2$) are those in Equation (27). In the lag columns, '0' entries denote that the lag is zero while '+' entries denote a positive value, and '-' entries denote a negative value (respectively greater or less relative to the base node's spatiotemporal position). Note that in case where one modality (space or time) shares the same (non-zero) sign, whether those two lags are identical or not qualitatively changes the spike sequence. The configuration column illustrates a schematic of the qualitative lag-motif, where the solid dot, small open circle, and large open circle indicate the base, second, and third nodes, respectively. The 14 motif classes are labelled by Roman numerals in the third column, information flow pattern.

| Lag-Sign Motif | $n_1$ | $t_1$ | $n_2$ | $t_2$ | Constraints | Motif Class | Configuration |
|---|---|---|---|---|---|---|---|
| 0 | 0 | 0 | 0 | 0 | | 0 | 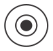 |
| 1.0 | 0 | 0 | 0 | + | | I | 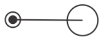 |
| 1.1 | 0 | 0 | 0 | - | | I | 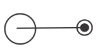 |
| 2.0 | 0 | 0 | + | 0 | | III | 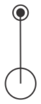 |

| | | | | | | | |
|---|---|---|---|---|---|---|---|
| 2.1 | 0 | 0 | - | 0 | | III | 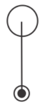 |
| 3.0 | 0 | 0 | + | + | | V | 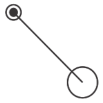 |
| 3.1 | 0 | 0 | + | - | | V | 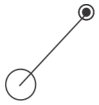 |
| 3.2 | 0 | 0 | - | + | | V | 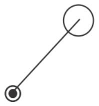 |
| 3.3 | 0 | 0 | - | - | | V | 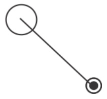 |
| 4.0 | 0 | + | 0 | 0 | | I | 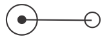 |
| 4.1 | 0 | - | 0 | 0 | | I | 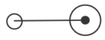 |
| 5.0.0 | 0 | + | 0 | + | $\lvert t_1 \rvert = \lvert t_2 \rvert$ | I | 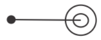 |
| 5.0.1 | 0 | + | 0 | + | $\lvert t_1 \rvert > \lvert t_2 \rvert$ | II | 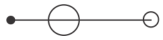 |
| 5.0.2 | 0 | + | 0 | + | $\lvert t_1 \rvert < \lvert t_2 \rvert$ | II | 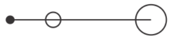 |
| 5.1 | 0 | + | 0 | - | | II | 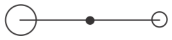 |
| 5.2 | 0 | - | 0 | + | | II | 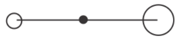 |
| 5.3.0 | 0 | - | 0 | - | $\lvert t_1 \rvert = \lvert t_2 \rvert$ | I | 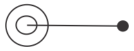 |
| 5.3.1 | 0 | - | 0 | - | $\lvert t_1 \rvert > \lvert t_2 \rvert$ | II | 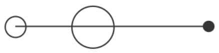 |
| 5.3.2 | 0 | - | 0 | - | $\lvert t_1 \rvert < \lvert t_2 \rvert$ | II | 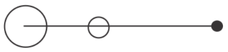 |
| 6.0 | 0 | + | + | 0 | | VI | 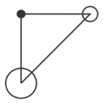 |
| 6.1 | 0 | + | - | 0 | | VI | 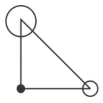 |

| | | | | | | | |
|---|---|---|---|---|---|---|---|
| 6.2 | 0 | - | + | 0 | | VII | 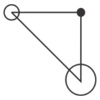 |
| 6.3 | 0 | - | - | 0 | | VII | 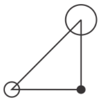 |
| 7.0.0 | 0 | + | + | + | $\lvert t_1\rvert = \lvert t_2\rvert$ | VII | 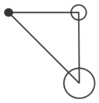 |
| 7.0.1 | 0 | + | + | + | $\lvert t_1\rvert > \lvert t_2\rvert$ | IX | 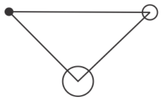 |
| 7.0.2 | 0 | + | + | + | $\lvert t_1\rvert < \lvert t_2\rvert$ | X | 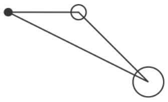 |
| 7.1 | 0 | + | + | - | | VIII | 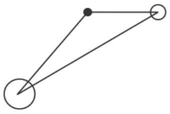 |
| 7.2.0 | 0 | + | - | + | $\lvert t_1\rvert = \lvert t_2\rvert$ | VII | 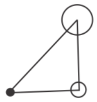 |
| 7.2.1 | 0 | + | - | + | $\lvert t_1\rvert > \lvert t_2\rvert$ | IX | 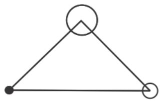 |
| 7.2.2 | 0 | + | - | + | $\lvert t_1\rvert < \lvert t_2\rvert$ | X | 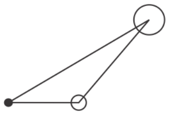 |
| 7.3 | 0 | + | - | - | | VIII | 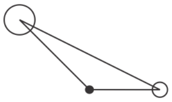 |
| 7.4 | 0 | - | + | + | | X | 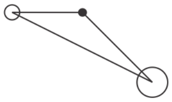 |
| 7.5.0 | 0 | - | + | - | $\lvert t_1\rvert = \lvert t_2\rvert$ | VI | 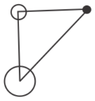 |

| | | | | | | | |
|---|---|---|---|---|---|---|---|
| 7.5.1 | 0 | - | + | - | $\lvert t_1 \rvert > \lvert t_2 \rvert$ | IX | 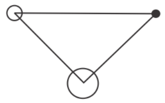 |
| 7.5.2 | 0 | - | + | - | $\lvert t_1 \rvert < \lvert t_2 \rvert$ | VIII | 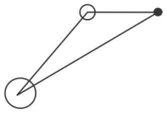 |
| 7.6 | 0 | - | - | + | | X | 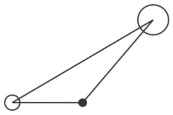 |
| 7.7.0 | 0 | - | - | - | $\lvert t_1 \rvert = \lvert t_2 \rvert$ | VI | 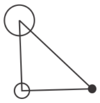 |
| 7.7.1 | 0 | - | - | - | $\lvert t_1 \rvert > \lvert t_2 \rvert$ | IX | 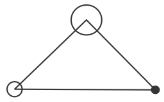 |
| 7.7.2 | 0 | - | - | - | $\lvert t_1 \rvert < \lvert t_2 \rvert$ | VIII | 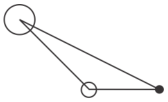 |
| 8.0 | + | 0 | 0 | 0 | | III | 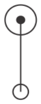 |
| 8.1 | - | 0 | 0 | 0 | | III | 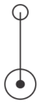 |
| 9.0 | + | 0 | 0 | + | | VI | 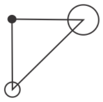 |
| 9.1 | + | 0 | 0 | - | | VII | 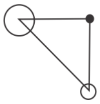 |
| 9.2 | - | 0 | 0 | + | | VI | 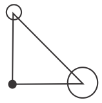 |
| 9.3 | - | 0 | 0 | - | | VII | 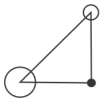 |

| | | | | | | | |
|---|---|---|---|---|---|---|---|
| 10.0.0 | + | 0 | + | 0 | $\lvert n_1 \rvert = \lvert n_2 \rvert$ | III | 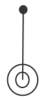 |
| 10.0.1 | + | 0 | + | 0 | $\lvert n_1 \rvert > \lvert n_2 \rvert$ | IV | 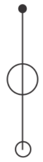 |
| 10.0.2 | + | 0 | + | 0 | $\lvert n_1 \rvert < \lvert n_2 \rvert$ | IV | 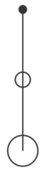 |
| 10.1 | + | 0 | - | 0 | | IV | 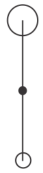 |
| 10.2 | - | 0 | + | 0 | | IV | 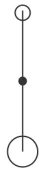 |
| 10.3.0 | - | 0 | - | 0 | $\lvert n_1 \rvert = \lvert n_2 \rvert$ | III | 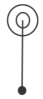 |
| 10.3.1 | - | 0 | - | 0 | $\lvert n_1 \rvert > \lvert n_2 \rvert$ | IV | 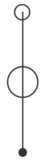 |
| 10.3.2 | - | 0 | - | 0 | $\lvert n_1 \rvert < \lvert n_2 \rvert$ | IV | 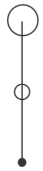 |
| 11.0.0 | + | 0 | + | + | $\lvert n_1 \rvert = \lvert n_2 \rvert$ | VI | 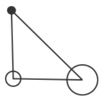 |

| | | | | | | | |
|---|---|---|---|---|---|---|---|
| 11.0.1 | + | 0 | + | + | $\lvert n_1\rvert > \lvert n_2\rvert$ | XII | 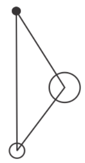 |
| 11.0.2 | + | 0 | + | + | $\lvert n_1\rvert < \lvert n_2\rvert$ | XII | 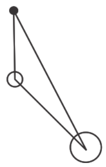 |
| 11.1.0 | + | 0 | + | - | $\lvert n_1\rvert = \lvert n_2\rvert$ | VII | 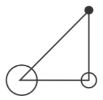 |
| 11.1.1 | + | 0 | + | - | $\lvert n_1\rvert > \lvert n_2\rvert$ | XI | 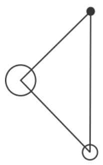 |
| 11.1.2 | + | 0 | + | - | $\lvert n_1\rvert < \lvert n_2\rvert$ | XI | 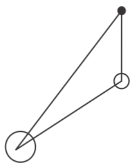 |
| 11.2 | + | 0 | - | + | | XII | 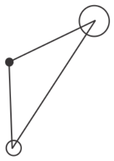 |
| 11.3 | + | 0 | - | - | | XI | 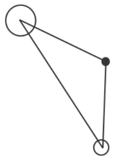 |
| 11.4 | - | 0 | + | + | | XII | 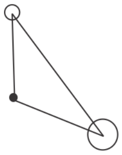 |
| 11.5 | - | 0 | + | - | | XI | 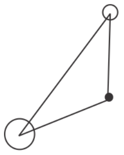 |

| | | | | | | | |
|---|---|---|---|---|---|---|---|
| 11.6.0 | - | 0 | - | + | $\|n_1\| = \|n_2\|$ | VI | 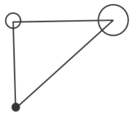 |
| 11.6.1 | - | 0 | - | + | $\|n_1\| > \|n_2\|$ | XII | 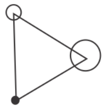 |
| 11.6.2 | - | 0 | - | + | $\|n_1\| < \|n_2\|$ | XII | 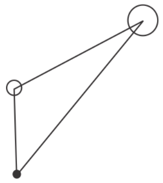 |
| 11.7.0 | - | 0 | - | - | $\|n_1\| = \|n_2\|$ | VII | 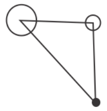 |
| 11.7.1 | - | 0 | - | - | $\|n_1\| > \|n_2\|$ | XI | 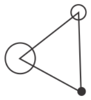 |
| 11.7.2 | - | 0 | - | - | $\|n_1\| < \|n_2\|$ | XI | 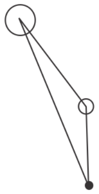 |
| 12.0 | + | + | 0 | 0 | | V | 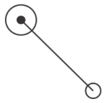 |
| 12.1 | + | - | 0 | 0 | | V | 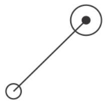 |
| 12.2 | - | + | 0 | 0 | | V | 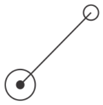 |
| 12.3 | - | - | 0 | 0 | | V | 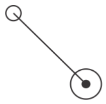 |
| 13.0.0 | + | + | 0 | + | $\|t_1\| = \|t_2\|$ | VII | 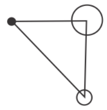 |

| | | | | | | | |
|---|---|---|---|---|---|---|---|
| 13.0.1 | + | + | 0 | + | $\lvert t_1 \rvert > \lvert t_2 \rvert$ | X | 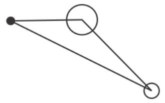 |
| 13.0.2 | + | + | 0 | + | $\lvert t_1 \rvert < \lvert t_2 \rvert$ | IX | 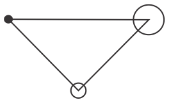 |
| 13.1 | + | + | 0 | - | | X | 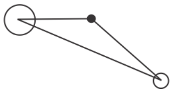 |
| 13.2 | + | - | 0 | + | | VIII | 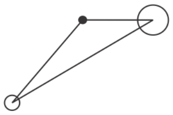 |
| 13.3.0 | + | - | 0 | - | $\lvert t_1 \rvert = \lvert t_2 \rvert$ | VI | 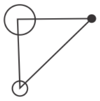 |
| 13.3.1 | + | - | 0 | - | $\lvert t_1 \rvert > \lvert t_2 \rvert$ | VIII | 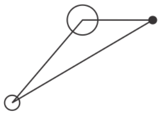 |
| 13.3.2 | + | - | 0 | - | $\lvert t_1 \rvert < \lvert t_2 \rvert$ | IX | 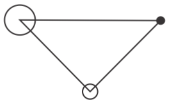 |
| 13.4.0 | - | + | 0 | + | $\lvert t_1 \rvert = \lvert t_2 \rvert$ | VII | 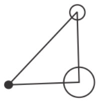 |
| 13.4.1 | - | + | 0 | + | $\lvert t_1 \rvert > \lvert t_2 \rvert$ | X | 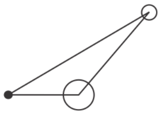 |
| 13.4.2 | - | + | 0 | + | $\lvert t_1 \rvert < \lvert t_2 \rvert$ | IX | 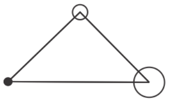 |
| 13.5 | - | + | 0 | - | | X | 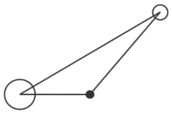 |
| 13.6 | - | - | 0 | + | | VIII | 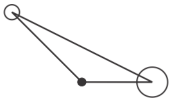 |

| | | | | | | | |
|---|---|---|---|---|---|---|---|
| 13.7.0 | - | - | 0 | - | $\vert\mathfrak{t}_1\vert = \vert\mathfrak{t}_2\vert$ | VI | 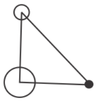 |
| 13.7.1 | - | - | 0 | - | $\vert\mathfrak{t}_1\vert > \vert\mathfrak{t}_2\vert$ | VIII | 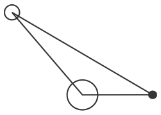 |
| 13.7.2 | - | - | 0 | - | $\vert\mathfrak{t}_1\vert < \vert\mathfrak{t}_2\vert$ | IX | 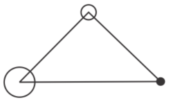 |
| 14.0.0 | + | + | + | 0 | $\vert\mathfrak{n}_1\vert = \vert\mathfrak{n}_2\vert$ | VI | 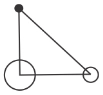 |
| 14.0.1 | + | + | + | 0 | $\vert\mathfrak{n}_1\vert > \vert\mathfrak{n}_2\vert$ | XII | 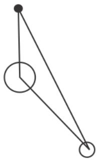 |
| 14.0.2 | + | + | + | 0 | $\vert\mathfrak{n}_1\vert < \vert\mathfrak{n}_2\vert$ | XII | 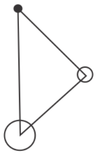 |
| 14.1 | + | + | - | 0 | | XII | 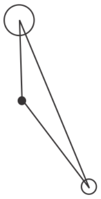 |
| 14.2.0 | + | - | + | 0 | $\vert\mathfrak{n}_1\vert = \vert\mathfrak{n}_2\vert$ | VII | 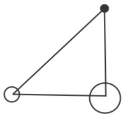 |
| 14.2.1 | + | - | + | 0 | $\vert\mathfrak{n}_1\vert > \vert\mathfrak{n}_2\vert$ | XI | 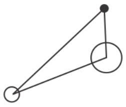 |
| 14.2.2 | + | - | + | 0 | $\vert\mathfrak{n}_1\vert < \vert\mathfrak{n}_2\vert$ | XI | 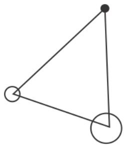 |

| | | | | | | | |
|---|---|---|---|---|---|---|---|
| 14.3 | + | - | - | 0 | | XI | 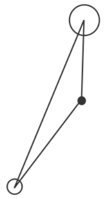 |
| 14.4 | - | + | + | 0 | | XII | 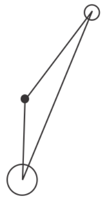 |
| 14.5.0 | - | + | - | 0 | $\lvert n_1 \rvert = \lvert n_2 \rvert$ | VI | 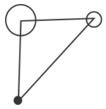 |
| 14.5.1 | - | + | - | 0 | $\lvert n_1 \rvert > \lvert n_2 \rvert$ | XII | 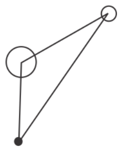 |
| 14.5.2 | - | + | - | 0 | $\lvert n_1 \rvert < \lvert n_2 \rvert$ | XII | 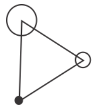 |
| 14.6 | - | - | + | 0 | | XI | 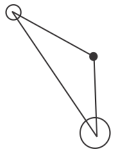 |
| 14.7.0 | - | - | - | 0 | $\lvert n_1 \rvert = \lvert n_2 \rvert$ | VII | 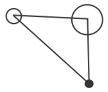 |
| 14.7.1 | - | - | - | 0 | $\lvert n_1 \rvert > \lvert n_2 \rvert$ | XI | 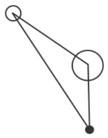 |
| 14.7.2 | - | - | - | 0 | $\lvert n_1 \rvert < \lvert n_2 \rvert$ | XI | 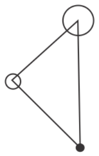 |

| | | | | | | | |
|---|---|---|---|---|---|---|---|
| 15.0.0 | + | + | + | + | $\lvert n_1 \rvert = \lvert n_2 \rvert$<br>$\lvert t_1 \rvert = \lvert t_2 \rvert$ | V | 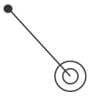 |
| 15.0.1 | + | + | + | + | $\lvert n_1 \rvert = \lvert n_2 \rvert$<br>$\lvert t_1 \rvert > \lvert t_2 \rvert$ | VIII | 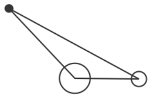 |
| 15.0.2 | + | + | + | + | $\lvert n_1 \rvert = \lvert n_2 \rvert$<br>$\lvert t_1 \rvert < \lvert t_2 \rvert$ | VIII | 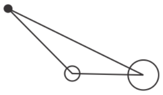 |
| 15.0.3 | + | + | + | + | $\lvert n_1 \rvert > \lvert n_2 \rvert$<br>$\lvert t_1 \rvert = \lvert t_2 \rvert$ | XI | 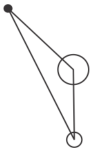 |
| 15.0.4 | + | + | + | + | $\lvert n_1 \rvert > \lvert n_2 \rvert$<br>$\lvert t_1 \rvert > \lvert t_2 \rvert$ | XIII | 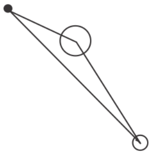 |
| 15.0.5 | + | + | + | + | $\lvert n_1 \rvert > \lvert n_2 \rvert$<br>$\lvert t_1 \rvert < \lvert t_2 \rvert$ | XIII | 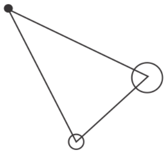 |
| 15.0.6 | + | + | + | + | $\lvert n_1 \rvert < \lvert n_2 \rvert$<br>$\lvert t_1 \rvert = \lvert t_2 \rvert$ | XI | 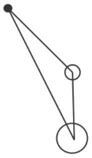 |
| 15.0.7 | + | + | + | + | $\lvert n_1 \rvert < \lvert n_2 \rvert$<br>$\lvert t_1 \rvert > \lvert t_2 \rvert$ | XIII | 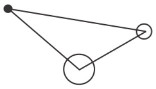 |
| 15.0.8 | + | + | + | + | $\lvert n_1 \rvert < \lvert n_2 \rvert$<br>$\lvert t_1 \rvert < \lvert t_2 \rvert$ | XIII | 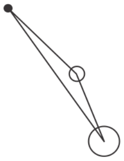 |
| 15.1.0 | + | + | + | - | $\lvert n_1 \rvert = \lvert n_2 \rvert$ | IX | 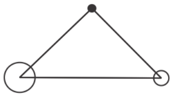 |

| | | | | | | | |
|---|---|---|---|---|---|---|---|
| 15.1.1 | + | + | + | - | $\|n_1\| > \|n_2\|$ | XIII | 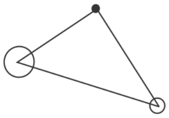 |
| 15.1.2 | + | + | + | - | $\|n_1\| < \|n_2\|$ | XIII | 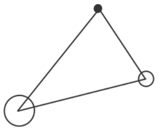 |
| 15.2.0 | + | + | - | + | $\|t_1\| = \|t_2\|$ | XI | 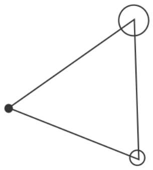 |
| 15.2.1 | + | + | - | + | $\|t_1\| > \|t_2\|$ | XIII | 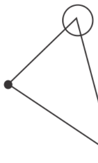 |
| 15.2.2 | + | + | - | + | $\|t_1\| < \|t_2\|$ | XIII | 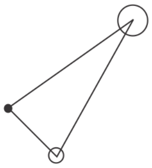 |
| 15.3 | + | + | - | - | | XIII | 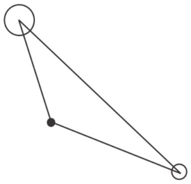 |
| 15.4.0 | + | - | + | + | $\|n_1\| = \|n_2\|$ | IX | 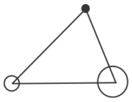 |
| 15.4.1 | + | - | + | + | $\|n_1\| > \|n_2\|$ | XIII | 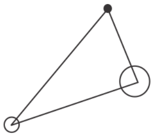 |
| 15.4.2 | + | - | + | + | $\|n_1\| < \|n_2\|$ | XIII | 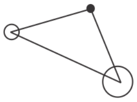 |

| | | | | | | | |
|---|---|---|---|---|---|---|---|
| 15.5.0 | + | - | + | - | $\lvert n_1\rvert = \lvert n_2\rvert$<br>$\lvert t_1\rvert = \lvert t_2\rvert$ | V | 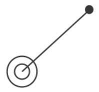 |
| 15.5.1 | + | - | + | - | $\lvert n_1\rvert = \lvert n_2\rvert$<br>$\lvert t_1\rvert > \lvert t_2\rvert$ | X | 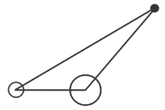 |
| 15.5.2 | + | - | + | - | $\lvert n_1\rvert = \lvert n_2\rvert$<br>$\lvert t_1\rvert < \lvert t_2\rvert$ | X | 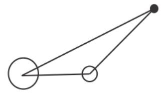 |
| 15.5.3 | + | - | + | - | $\lvert n_1\rvert > \lvert n_2\rvert$<br>$\lvert t_1\rvert = \lvert t_2\rvert$ | XII | 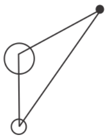 |
| 15.5.4 | + | - | + | - | $\lvert n_1\rvert > \lvert n_2\rvert$<br>$\lvert t_1\rvert > \lvert t_2\rvert$ | XIII | 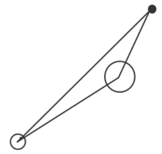 |
| 15.5.5 | + | - | + | - | $\lvert n_1\rvert > \lvert n_2\rvert$<br>$\lvert t_1\rvert < \lvert t_2\rvert$ | XIII | 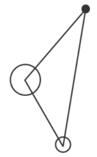 |
| 15.5.6 | + | - | + | - | $\lvert n_1\rvert < \lvert n_2\rvert$<br>$\lvert t_1\rvert = \lvert t_2\rvert$ | XII | 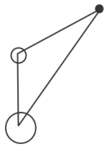 |
| 15.5.7 | + | - | + | - | $\lvert n_1\rvert < \lvert n_2\rvert$<br>$\lvert t_1\rvert > \lvert t_2\rvert$ | XIII | 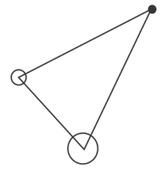 |
| 15.5.8 | + | - | + | - | $\lvert n_1\rvert < \lvert n_2\rvert$<br>$\lvert t_1\rvert < \lvert t_2\rvert$ | XIII | 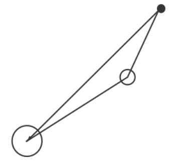 |

| | | | | | | | |
|---|---|---|---|---|---|---|---|
| 15.6 | + | - | - | + | | XIII | 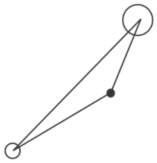 |
| 15.7.0 | + | - | - | - | $\lvert t_1 \rvert = \lvert t_2 \rvert$ | XII | 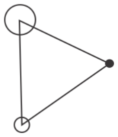 |
| 15.7.1 | + | - | - | - | $\lvert t_1 \rvert > \lvert t_2 \rvert$ | XIII | 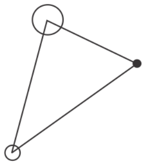 |
| 15.7.2 | + | - | - | - | $\lvert t_1 \rvert < \lvert t_2 \rvert$ | XIII | 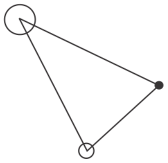 |
| 15.8.0 | - | + | + | + | $\lvert t_1 \rvert = \lvert t_2 \rvert$ | XI | 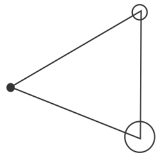 |
| 15.8.1 | - | + | + | + | $\lvert t_1 \rvert > \lvert t_2 \rvert$ | XIII | 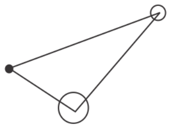 |
| 15.8.2 | - | + | + | + | $\lvert t_1 \rvert < \lvert t_2 \rvert$ | XIII | 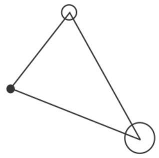 |
| 15.9 | - | + | + | - | | XIII | 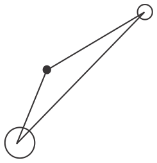 |
| 15.10.0 | - | + | - | + | $\lvert n_1 \rvert = \lvert n_2 \rvert$ <br> $\lvert t_1 \rvert = \lvert t_2 \rvert$ | V | 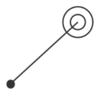 |

| | | | | | | | |
|---|---|---|---|---|---|---|---|
| 15.10.1 | - | + | - | + | $\lvert\mathfrak{n}_1\rvert = \lvert\mathfrak{n}_2\rvert$<br>$\lvert\mathfrak{t}_1\rvert > \lvert\mathfrak{t}_2\rvert$ | VIII | 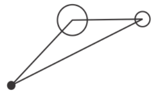 |
| 15.10.2 | - | + | - | + | $\lvert\mathfrak{n}_1\rvert = \lvert\mathfrak{n}_2\rvert$<br>$\lvert\mathfrak{t}_1\rvert < \lvert\mathfrak{t}_2\rvert$ | VIII | 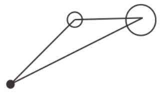 |
| 15.10.3 | - | + | - | + | $\lvert\mathfrak{n}_1\rvert > \lvert\mathfrak{n}_2\rvert$<br>$\lvert\mathfrak{t}_1\rvert = \lvert\mathfrak{t}_2\rvert$ | XI | 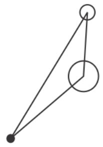 |
| 15.10.4 | - | + | - | + | $\lvert\mathfrak{n}_1\rvert > \lvert\mathfrak{n}_2\rvert$<br>$\lvert\mathfrak{t}_1\rvert > \lvert\mathfrak{t}_2\rvert$ | XIII | 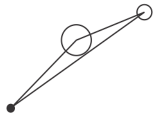 |
| 15.10.5 | - | + | - | + | $\lvert\mathfrak{n}_1\rvert > \lvert\mathfrak{n}_2\rvert$<br>$\lvert\mathfrak{t}_1\rvert < \lvert\mathfrak{t}_2\rvert$ | XIII | 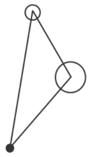 |
| 15.10.6 | - | + | - | + | $\lvert\mathfrak{n}_1\rvert < \lvert\mathfrak{n}_2\rvert$<br>$\lvert\mathfrak{t}_1\rvert = \lvert\mathfrak{t}_2\rvert$ | XI | 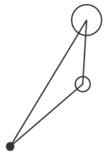 |
| 15.10.7 | - | + | - | + | $\lvert\mathfrak{n}_1\rvert < \lvert\mathfrak{n}_2\rvert$<br>$\lvert\mathfrak{t}_1\rvert > \lvert\mathfrak{t}_2\rvert$ | XIII | 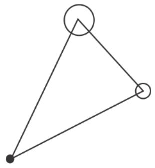 |
| 15.10.8 | - | + | - | + | $\lvert\mathfrak{n}_1\rvert < \lvert\mathfrak{n}_2\rvert$<br>$\lvert\mathfrak{t}_1\rvert < \lvert\mathfrak{t}_2\rvert$ | XIII | 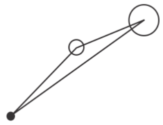 |
| 15.11.0 | - | + | - | - | $\lvert\mathfrak{n}_1\rvert = \lvert\mathfrak{n}_2\rvert$ | IX | 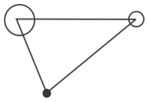 |
| 15.11.1 | - | + | - | - | $\lvert\mathfrak{n}_1\rvert > \lvert\mathfrak{n}_2\rvert$ | XIII | 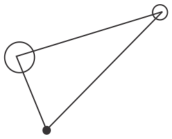 |

| | | | | | | | |
|---|---|---|---|---|---|---|---|
| 15.11.2 | - | + | - | - | $\lvert n_1\rvert < \lvert n_2\rvert$ | XIII | 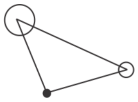 |
| 15.12 | - | - | + | + | | XIII | 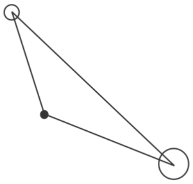 |
| 15.13.0 | - | - | + | - | $\lvert t_1\rvert = \lvert t_2\rvert$ | XII | 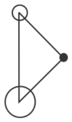 |
| 15.13.1 | - | - | + | - | $\lvert t_1\rvert > \lvert t_2\rvert$ | XIII | 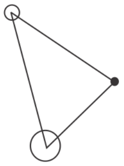 |
| 15.13.2 | - | - | + | - | $\lvert t_1\rvert < \lvert t_2\rvert$ | XIII | 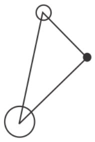 |
| 15.14.0 | - | - | - | + | $\lvert n_1\rvert = \lvert n_2\rvert$ | IX | 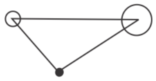 |
| 15.14.1 | - | - | - | + | $\lvert n_1\rvert > \lvert n_2\rvert$ | XIII | 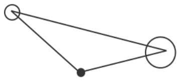 |
| 15.14.2 | - | - | - | + | $\lvert n_1\rvert < \lvert n_2\rvert$ | XIII | 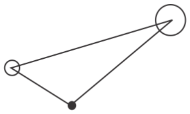 |
| 15.15.0 | - | - | - | - | $\lvert n_1\rvert = \lvert n_2\rvert$ <br> $\lvert t_1\rvert = \lvert t_2\rvert$ | V | 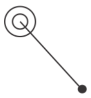 |
| 15.15.1 | - | - | - | - | $\lvert n_1\rvert = \lvert n_2\rvert$ <br> $\lvert t_1\rvert > \lvert t_2\rvert$ | X | 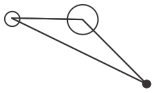 |
| 15.15.2 | - | - | - | - | $\lvert n_1\rvert = \lvert n_2\rvert$ <br> $\lvert t_1\rvert < \lvert t_2\rvert$ | X | 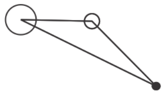 |

| | | | | | | | |
|---|---|---|---|---|---|---|---|
| 15.15.3 | - | - | - | - | $\lvert n_1 \rvert > \lvert n_2 \rvert$<br>$\lvert t_1 \rvert = \lvert t_2 \rvert$ | XII | 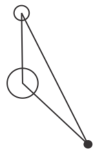 |
| 15.15.4 | - | - | - | - | $\lvert n_1 \rvert > \lvert n_2 \rvert$<br>$\lvert t_1 \rvert > \lvert t_2 \rvert$ | XIII | 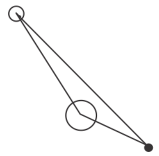 |
| 15.15.5 | - | - | - | - | $\lvert n_1 \rvert > \lvert n_2 \rvert$<br>$\lvert t_1 \rvert < \lvert t_2 \rvert$ | XIII | 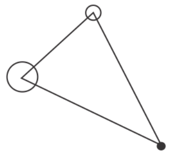 |
| 15.15.6 | - | - | - | - | $\lvert n_1 \rvert < \lvert n_2 \rvert$<br>$\lvert t_1 \rvert = \lvert t_2 \rvert$ | XII | 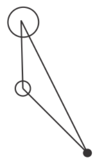 |
| 15.15.7 | - | - | - | - | $\lvert n_1 \rvert < \lvert n_2 \rvert$<br>$\lvert t_1 \rvert > \lvert t_2 \rvert$ | XIII | 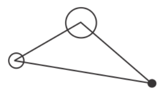 |
| 15.15.8 | - | - | - | - | $\lvert n_1 \rvert < \lvert n_2 \rvert$<br>$\lvert t_1 \rvert < \lvert t_2 \rvert$ | XIII | 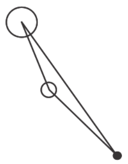 |

2. Dechery, J. B. & MacLean, J. N. Functional triplet motifs underlie accurate predictions of single-trial responses in populations of tuned and untuned V1 neurons. *PLOS Computational Biology* **14**, e1006153, DOI: 10.1371/journal.pcbi.1006153

3. Schneidman, E., Berry, M. J., Segev, R. & Bialek, W. Weak pairwise correlations imply strongly correlated network states in a neural population. *Nature* **440**, 1007–1012, DOI: 10.1038/nature04701

4. Jiang, M., Gao, X., An, H., Li, H. & Sun, B. Reconstructing complex network for characterizing the time-varying causality evolution behavior of multivariate time series. *Scientific Reports* **7**, 10486, DOI: 10.1038/s41598-017-10759-3

5. Sporns, O. & Kötter, R. Motifs in Brain Networks. *PLoS Biology* **2**, e369, DOI: 10.1371/journal.pbio.0020369

6. Jovanović, S. & Rotter, S. Interplay between Graph Topology and Correlations of Third Order in Spiking Neuronal Networks. *PLOS Computational Biology* **12**, e1004963, DOI: 10.1371/journal.pcbi.1004963

7. Milo, R. *et al.* Network Motifs: Simple Building Blocks of Complex Networks. *Science* **298**, 824–827, DOI: 10.1126/science.298.5594.824

8. Buzsáki, G. & Draguhn, A. Neuronal oscillations in cortical networks. *Science* **304**, 1926–1929, DOI: 10.1126/science.1099745

9. Yu, S. *et al.* Higher-Order Interactions Characterized in Cortical Activity. *Journal of Neuroscience* **31**, 17514–17526, DOI: 10.1523/JNEUROSCI.3127-11.2011

10. Bartelt, H., Lohmann, A. W. & Wirnitzer, B. Phase and amplitude recovery from bispectra. *Applied Optics* **23**, 3121–3129, DOI: 10.1364/AO.23.003121

11. Julesz, B. Visual Pattern Discrimination. *IRE Transactions on Information Theory* **8**, 84–92, DOI: 10.1109/TIT.1962.1057698

12. Julesz, B., Gilbert, E. N., Shepp, L. A. & Frisch, H. L. Inability of humans to discriminate between visual textures that agree in second-order statistics-revisited. *Perception* **2**, 391–405, DOI: 10.1068/p020391

13. Yellott, J. I. & Iverson, G. J. Uniqueness properties of higher-order autocorrelation functions. *Journal of the Optical Society of America A* **9**, 388–404, DOI: 10.1364/JOSAA.9.000388

14. Yellott, J. I. Implications of triple correlation uniqueness for texture statistics and the Julesz conjecture. *Journal of the Optical Society of America A* **10**, 777–793, DOI: 10.1364/JOSAA.10.000777

15. Victor, J. D. Images, statistics, and textures: implications of triple correlation uniqueness for texture statistics and the Julesz conjecture: comment. *J. Opt. Soc. Am. A, JOSAA* **11**, 1680–1684, DOI: 10.1364/JOSAA.11.001680

16. Feller, W. *An introduction to probability theory and its applications* (Wiley, Oxford, 1950),

17. Roscoe, D. D., Hamm, T. M., Reinking, R. M. & Stuart, D. G. Detection of synchrony in the discharge of a population of neurons. II. Implementation and sensitivity of a synchronization index. *J Neurosci Methods* **13**, 51–64, DOI: 10.1016/0165-0270(85)90043-3

18. Kreuz, T., Chicharro, D., Greschner, M. & Andrzejak, R. G. Time-resolved and time-scale adaptive measures of spike train synchrony. *J Neurosci Methods* **195**, 92–106, DOI: 10.1016/j.jneumeth.2010.11.020

19. Kreuz, T., Chicharro, D., Houghton, C., Andrzejak, R. G. & Mormann, F. Monitoring spike train synchrony. *Journal of Neurophysiology* **109**, 1457–1472, DOI: 10.1152/jn.00873.2012

20. Hyvärinen, T. *et al.* Functional characterization of human pluripotent stem cell-derived cortical networks differentiated on laminin-521 substrate: comparison to rat cortical cultures. *Sci Rep* **9**, 17125, DOI: 10.1038/s41598-019-53647-8

21. Kapucu, F. E., Vinogradov, A., Hyvärinen, T., Ylä-Outinen, L. & Narkilahti, S. Comparative microelectrode array data of the functional development of hPSC-derived and rat neuronal networks. *Scientific Data* **9**, 1–10

22. Suresh, J. *et al.* Network burst activity in hippocampal neuronal cultures: the role of synaptic and intrinsic currents. *Journal of Neurophysiology* **115**, 3073–3089

23. McCulloch, W. S. & Pitts, W. A logical calculus of the ideas immanent in nervous activity. *Bulletin of Mathematical Biophysics* **5**, 115–133, DOI: 10.1007/BF02478259

24. Hebb, D. O. *The organization of behavior; a neuropsychological theory* (Wiley, Oxford, England, 1949)

25. Abeles, M. *Corticonics: Neural Circuits of the Cerebral Cortex* (Cambridge University Press, Cambridge, 1991)

26. Singer, W. Neuronal Synchrony: A Versatile Code for the Definition of Relations? *Neuron* **24**, 49–65, DOI: 10.1016/S0896-6273(00)80821-1

27. Russo, E. & Durstewitz, D. Cell assemblies at multiple time scales with arbitrary lag constellations. *eLife* **6**, e19428, DOI: 10.7554/eLife.19428

28. Skaggs, W. E. & McNaughton, B. L. Replay of neuronal firing sequences in rat hippocampus during sleep following spatial experience. *Science* **271**, 1870–1873, DOI: 10.1126/science.271.5257.1870

29. Abeles, M. *Local Cortical Circuits*, vol. 6 of *Studies of Brain Function* (Springer, Berlin, 1982),

30. Hertz, J. & Prügel-Bennett, A. Learning synfire chains: turning noise into signal. *Int J Neural Syst* **7**, 445–450, DOI: 10.1142/s0129065796000427

31. Diesmann, M., Gewaltig, M.-O. & Aertsen, A. Stable propagation of synchronous spiking in cortical neural networks. *Nature* **402**, 529–533, DOI: 10.1038/990101

32. Ikegaya, Y. *et al.* Synfire chains and cortical songs: temporal modules of cortical activity. *Science* **304**, 559–564, DOI: 10.1126/science.1093173

33. Jirsa, V. K., Stacey, W. C., Quilichini, P. P., Ivanov, A. I. & Bernard, C. On the nature of seizure dynamics. *Brain* **137**, 2210–2230, DOI: 10.1093/brain/awu133

34. Riehle, A., Grün, S., Diesmann, M. & Aertsen, A. Spike synchronization and rate modulation differentially involved in motor cortical function. *Science* **278**, 1950–1953, DOI: 10.1126/science.278.5345.1950

35. Saper, C. B., Fuller, P. M., Pedersen, N. P., Lu, J. & Scammell, T. E. Sleep State Switching. *Neuron* **68**, 1023–1042, DOI: 10.1016/j.neuron.2010.11.032

36. Iorio, F., Saez-Rodriguez, J. & Bernardo, D. d. Network based elucidation of drug response: from modulators to targets. *BMC Syst Biol* **7**, 139, DOI: 10.1186/1752-0509-7-139

37. Stella, A., Bouss, P., Palm, G. & Grün, S. Comparing Surrogates to Evaluate Precisely Timed Higher-Order Spike Correlations. *eNeuro* **9**, DOI: 10.1523/ENEURO.0505-21.2022

38. Baker, S. N. & Lemon, R. N. Precise spatiotemporal repeating patterns in monkey primary and supplementary motor areas occur at chance levels. *Journal of neurophysiology* **84**, 1770–1780

39. Grün, S. Data-Driven Significance Estimation for Precise Spike Correlation. *Journal of Neurophysiology* **101**, 1126–1140, DOI: 10.1152/jn.00093.2008

40. van Drongelen, W. *Signal processing for neuroscientists: Introduction to the analysis of physiological signals.* (Elsevier/Academic Press, 2018), Chapter 7

41. Bojanek, K., Zhu, Y. & MacLean, J. Cyclic transitions between higher order motifs underlie sustained asynchronous spiking in sparse recurrent networks. *PLoS computational biology* **16**, e1007409

42. Bou Assi, E., Gagliano, L., Rihana, S., Nguyen, D. K. & Sawan, M. Bispectrum Features and Multilayer Perceptron Classifier to Enhance Seizure Prediction. *Sci Rep* **8**, 15491, DOI: 10.1038/s41598-018-33969-9

43. Gagliano, L., Bou Assi, E., Nguyen, D. K. & Sawan, M. Bispectrum and Recurrent Neural Networks: Improved Classification of Interictal and Preictal States. *Sci Rep* **9**, 15649, DOI: 10.1038/s41598-019-52152-2

44. Jansen, N. A. *et al.* Impaired β-γ Coupling Indicates Inhibitory Dysfunction and Seizure Risk in a Dravet Syndrome Mouse Model. *The Journal of Neuroscience: The Official Journal of the Society for Neuroscience* **41**, 524–537, DOI: 10.1523/JNEUROSCI.2132-20.2020

45. Z. Moszner. Sur l'équation f(x)f(y)f(−x−y)=g(x)g(y)g(−x−y). *aequationes mathematicae* **21**, 20–32 (1980). doi:10.1007/BF02189336.

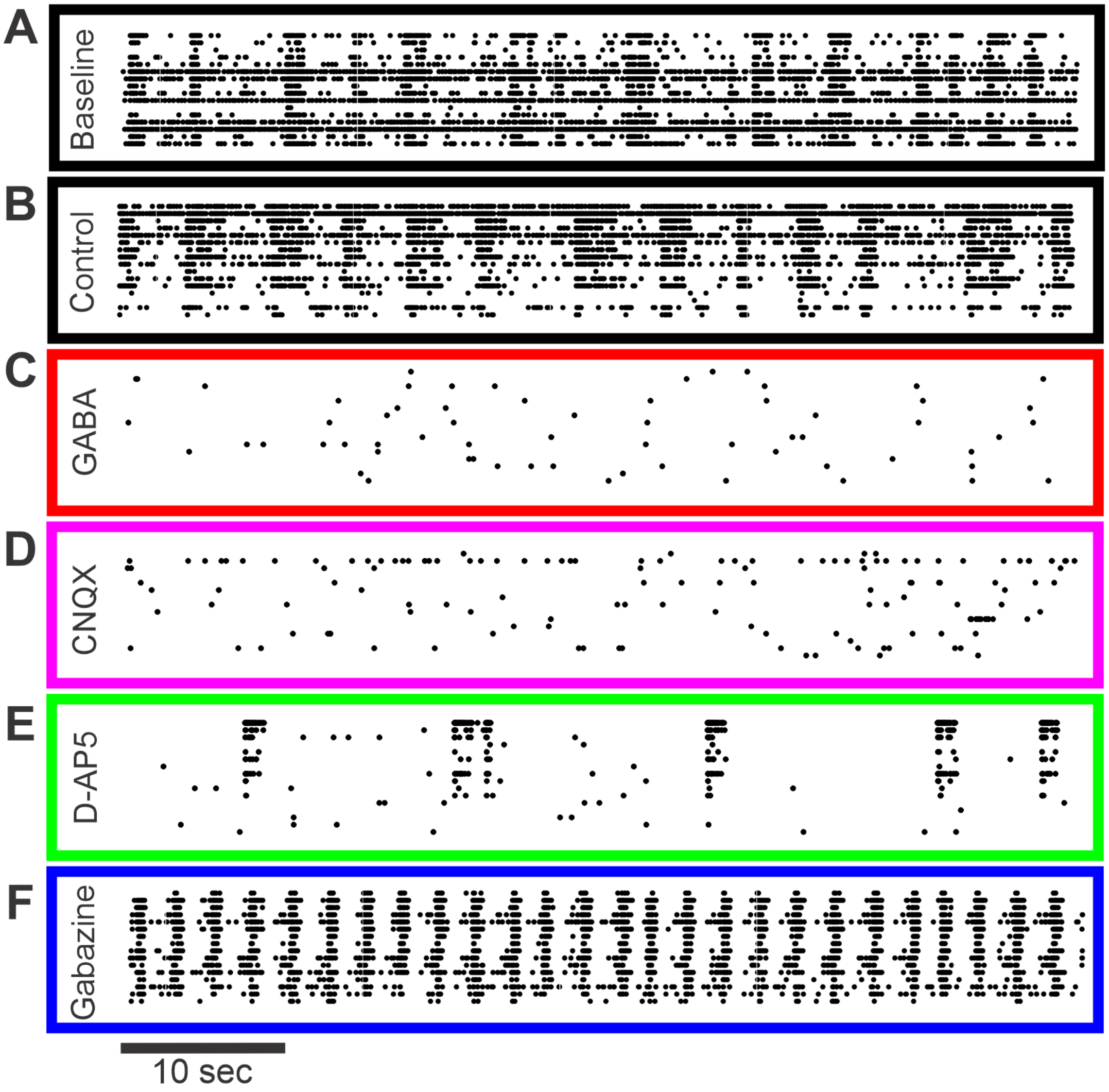

Figure S1: **Representative 1-minute epochs of rat cortical spike rasters per each experimental condition at 22 days in vitro [CITE Kapucu 2022].** Rat cortical cultures from microelectrode array well plates were exposed to the following pharmacological agents: baseline (A; black), control (B; black), GABA (C; red), CNQX (D; magenta), D-AP5 (E; green), and gabazine (F; blue). Note that we present the raster in two dimensions here while in our analysis for Fig. 5 we used the two spatial dimensions of the MEA and one temporal dimension.
